# Modeling the prodromal phase of Alzheimer’s disease: Selective amyloid-driven failure of cholinergic medial septal neurons perturbs REM sleep, cognition, emotion, and broadcasts pathology in aging mice

**DOI:** 10.1101/2025.07.09.663930

**Authors:** Mathieu Nollet, Wei Ba, Berta Anuncibay Soto, Chunyu Yin, Leda Lignos, Katarina Jovic, Sara Wong, Alexei L. Vyssotski, Raquel Yustos, Nicholas P. Franks, William Wisden

## Abstract

In humans, decreases in rapid eye movement sleep (REMS) strongly predict Alzheimer’s disease (AD), alongside early degeneration of basal forebrain cholinergic neurons. We examined how β-amyloid pathology gradually erodes mouse cholinergic neurons. The familial App^NL-G-F^ allele was selectively expressed in medial septal (MS) cholinergic neurons of both sexes and compared with mice with global App^NL-G-F^ expression and selective genetic lesions of MS cholinergic cells. By 14 months, targeted App^NL-G-F^ allele expression had caused loss of 25% of MS cholinergic neurons and produced amyloid deposition in their terminal fields, particularly the hippocampus. REMS was reduced, together with cognitive and emotional alterations mirroring phenotypes in global mutants, which also showed selective MS cholinergic cell loss. Genetic lesioning of MS cholinergic cells recapitulated most phenotypes, identifying cholinergic loss, not amyloid deposition, as a probable cause of these phenotypes. Nevertheless, broadcasted amyloid from MS cholinergic neurons likely induced hippocampal astrocyte activation and epileptiform spikes.

**Highlights:**

- Amyloid in medial septum causes cholinergic neuron loss and Alzheimer’s symptoms
- Amyloid from medial septal axons may trigger pathology in distant brain regions
- Loss of acetylcholine, not amyloid spread, drives REM sleep and cognitive deficits
- Broadcasted septal amyloid induces hippocampal astrogliosis and epileptiform spikes

**GRAPHICAL ABSTRACT:** 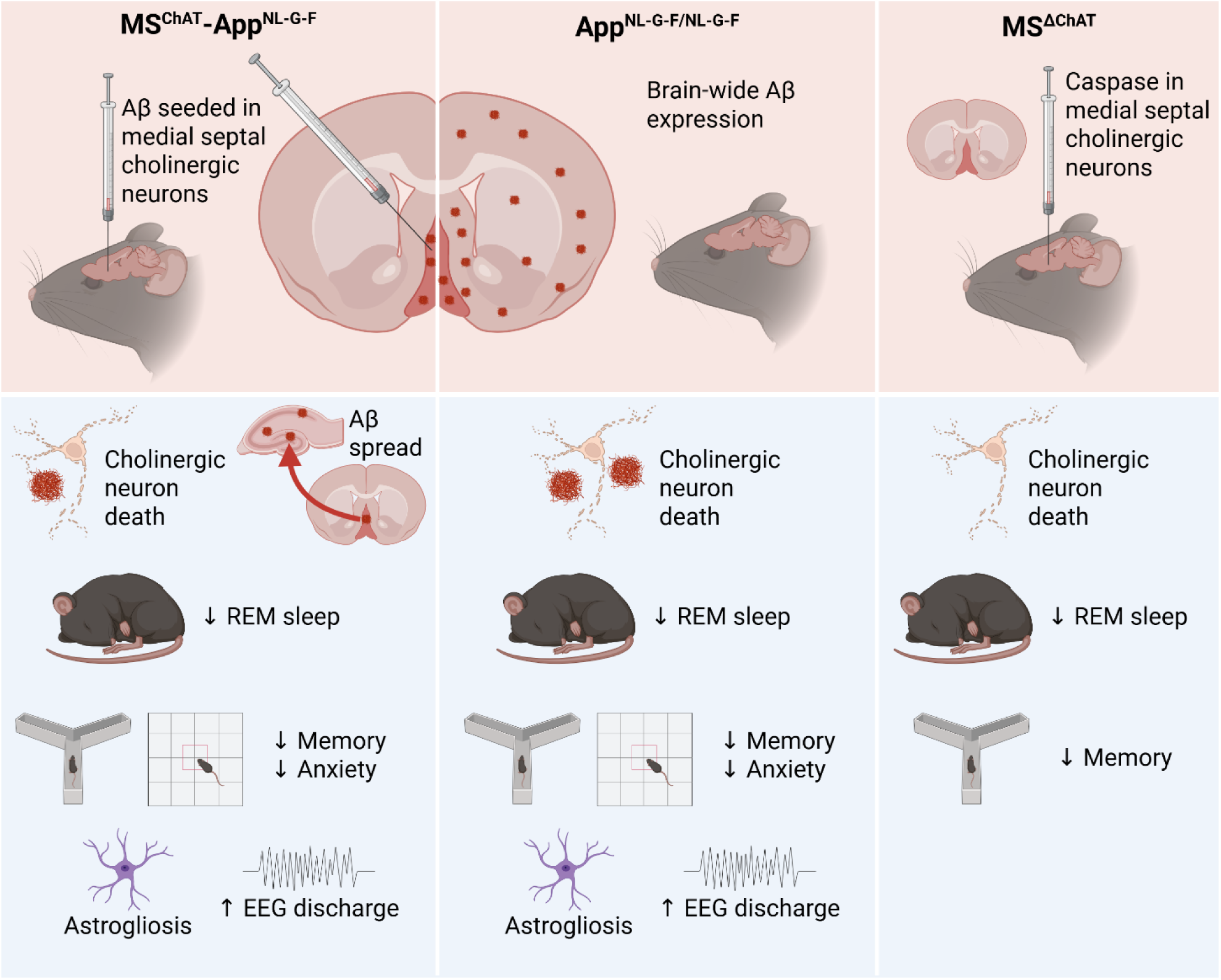

## INTRODUCTION

The fundamental nature of the AD disease(s) is conceived as a mixture of proteinopathies and inflammation, with age the biggest risk factor^1–4^. In the prodromal phase, the trajectory of the disease, however, may be shaped by selective declines in neuromodulator molecules that are produced subcortically^5^. For example, acetylcholine (ACh) is a neuromodulator, synthesized by the choline acetyltransferase (ChAT) enzyme, that supports arousal, emotion and cognition, activating both metabotropic (muscarinic) and ionotropic (nicotinic) receptors^6–15^. During wakefulness, but also during rapid eye movement sleep (REMS), ACh is released widely into the forebrain from key groups of neurons in the cholinergic basal forebrain (BF), which include, among other nuclei, the medial septum (MS)^6,7,16^.

Known for nearly half a century, and confirmed numerous times, the selective degeneration of BF-cholinergic neurons is a notable early feature of Alzheimer’s disease (AD)^17–30^, emerging before cortex atrophy^5,31,32^. It was postulated that this early decline of ACh neurons in humans causes the mild cognitive impairment at the start of AD^18^. Cholinergic esterase inhibitors significantly delay the decline of cognition for (some) patients^33^, but do not stop the disease progressing^24^. For these reasons, according to some commentators, interest in the cholinergic system’s failure in AD has languished^24,25^. It has also been suggested that neglecting the importance of neuromodulatory systems in the genesis of AD probably also reflects dominant cortico-centric models that assume that higher-order deficits are caused by cortical and hippocampal damage alone^13^. However, these cortico-centric models fail to account for the reliance of these structures on their neurochemical milieu and tend to minimize the critical role of ascending neuroregulatory systems in synchronizing flexible and precise information processing throughout the brain^13^.

Neuromodulators are also essential for orchestrating sleep-wake states^34,35^. Genetic knockout studies demonstrate that ACh is required for the generation of REMS, although the circuitry involved is unclear^36^. Importantly, reductions and alterations in REMS have emerged as strong predictors of clinical AD onset^37^, alongside alterations of non-rapid eye movement sleep (NREMS)^38^. Additionally, lower percentages of REMS correlate with smaller volumes of the cholinergic BF^39^ and are further linked with higher all-cause, cardiovascular, and non-cancer mortality^40^. Both REMS and its associated theta rhythms are hypothesized to promote cognition^41–43^, regulate emotion, and dampen stress responses^44–46^, processes that are compromised in AD and in individuals living with mild cognitive impairment, as well as in animal models exhibiting widespread amyloid-β (Aβ) accumulation^47–61^.

In humans, BF-cholinergic neurons are especially susceptible to Aβ toxicity, irrespective of the AD subtype, and BF atrophy correlates strongly with neocortical Aβ burden^21,62–64^. The degeneration of cholinergic neurons, alongside other neuromodulator systems, is posited to potentiate the pathological processes underlying AD^5,13^. Thus, it is worth examining the extent to which cholinergic deficits contribute to the disease symptoms. Of note, the MS contains a mixture of ACh, GABA and glutamate neurons which all project to the hippocampus and other areas^11,16,29,65–68^. Strikingly, Aβ peptide injection into the MS of rats disrupts neuronal activity, hippocampal long-term potentiation, theta rhythm and cognition^69,70^.

To test if and how selective amyloid pathology affects MS cholinergic (MS^ChAT^) cells, and its consequences for REMS and cognition, we expressed the synthetic *App*^NL-G-F^ allele, which comprises a combination of familial AD mutations^71^, selectively in mouse MS^ChAT^ neurons, and over 14 months, compared the phenotype in mice where the *App*^NL-G-F^ allele is expressed brain-wide. We found that inducing pathology specifically in the small population of MS^ChAT^ neurons generates highly similar phenotypes (*inter alia*, altered cognition and emotional processing, as well as selective REMS deficits) to those of the *App*^NL-G-F^ *knock-in* mouse^49,71^, where the allele is expressed in many cell types. Thus, the integrity of the medial septal cholinergic cells underpins a large part of *App*^NL-G-F^ model’s phenotypic features. In essence, we have proposed a circuit mechanism and developed an animal model that may capture key aspects of the prodromal phase of AD, which incorporates the selective vulnerability of acetylcholine cells and which suggests how an early and accumulating deficit in REM sleep and associated cognitive changes in the prodromal phases of AD might be produced. Nevertheless, broadcasted amyloid from MS cholinergic neurons also generates disease features as it produced hippocampal astrocyte activation and epileptiform spikes.

## RESULTS

### Cholinergic medial septal cells are wake- and REMS-active and primarily send projections to the hippocampus and cortex

We first characterized MS^ChAT^ cell activity across vigilance states. By calcium photometry, MS^ChAT^ cells were most active in REMS and WAKE, but quiet in NREMS (*AAV-DIO-GCaMP6s* was injected into the MS of *ChAT-Cre* mice; Figures 1A-B). Baseline Ca^2+^ transients increased during transitions from NREMS to REMS and NREMS to wakefulness. We next mapped the projection targets of MS^ChAT^ cells. Many of their axons were detected in hippocampal structures (CA areas, dentate gyrus and subiculum), but also in the medial prefrontal cortex (mPFC; prelimbic and cingulate cortices), primary cortices (somatosensory, motor and visual) and entorhinal cortex, as well as in the olfactory bulb, thalamus, hypothalamus, amygdala, and medial habenula (for the tracing, *AAV-DIO-ChR2-EYFP* was injected into the MS of *ChAT-Cre* mice; Figures 1C-D).

**Figure 1.**
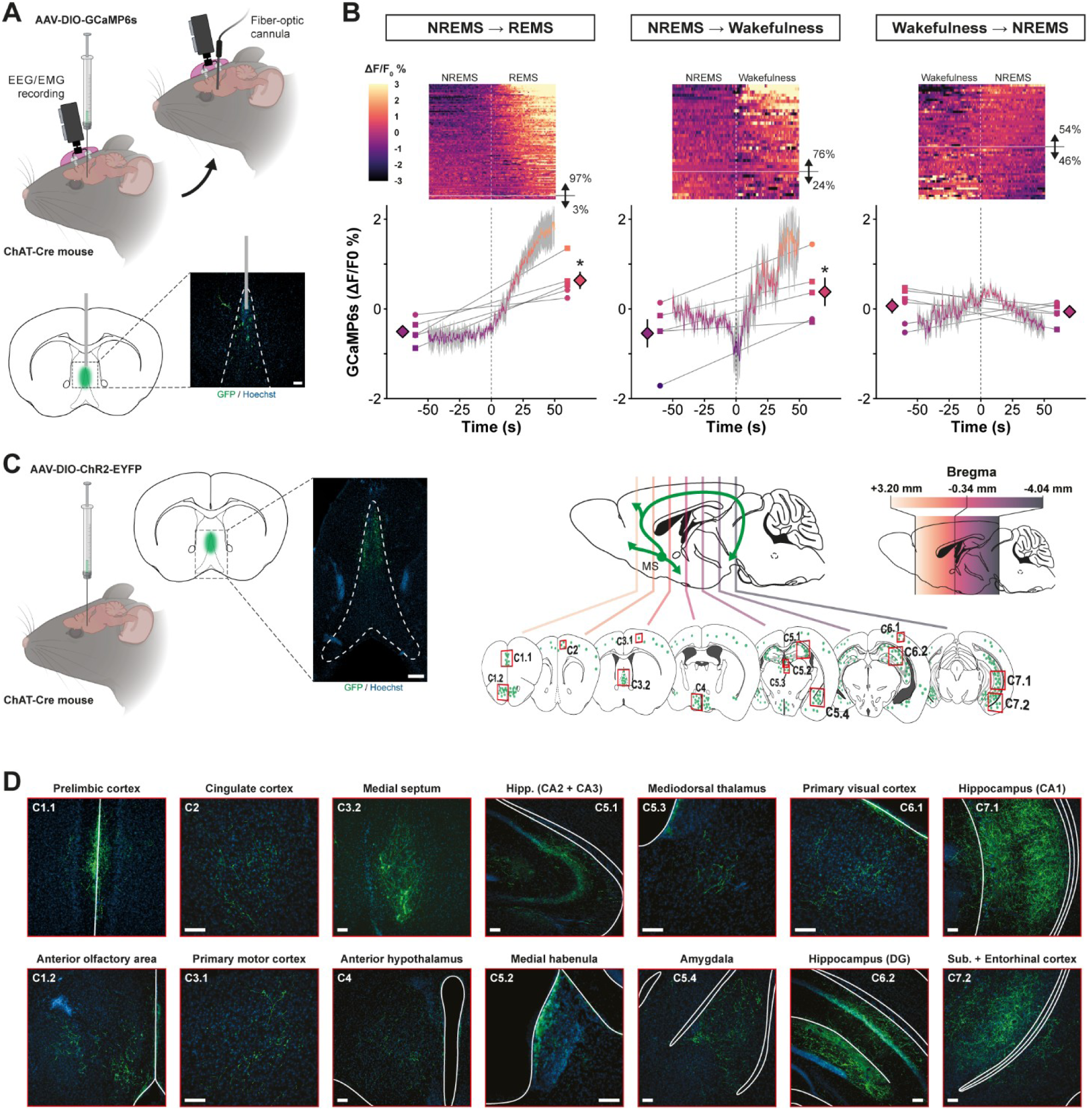
MS cholinergic neurons are active during REMS and wakefulness and predominantly project to hippocampal and cortical regions. (A) *AAV-DIO-GCaMP6s* was injected into the MS of *ChAT-Cre* mice to target cholinergic neurons (scale bar: 100 μm). An optic fiber was implanted above the MS, and EEG/EMG electrodes enabled simultaneous fiber photometry and polysomnographic recordings (*n* = 6 mice). (B) GCaMP fluorescence signals increased during transitions from NREMS to REMS and from NREMS to wakefulness, but showed minimal changes during wake-to-NREMS transitions, indicating selective activation of MS cholinergic neurons during REMS and wakefulness. During NREMS-to-REMS transitions, 97% of events were associated with increased activity, whereas 76% of NREMS-to-wakefulness transitions showed elevated firing. Transitions from wakefulness to NREMS displayed a balanced distribution of increases and decreases. (C) *AAV-DIO-ChR2-EYFP* was injected into the MS of *ChAT-Cre* mice (*n* = 2) to visualize cholinergic axons (green; scale bar: 200 μm). Serial coronal sections were used to map cholinergic projections. Gradient color-coding indicates the rostrocaudal position of each brain slice in whole-brain quantified immunohistochemical data. (D) Coronal brain sections show GFP-positive cholinergic projections (green) and Hoechst-stained nuclei (blue) including cortical, hippocampal, thalamic, and hypothalamic regions (scale bar: 100 μm). Crop identifiers (C1.1-C7.2) correspond to red squares in the schematic overview shown in panel C. Statistical analysis: two-sided Student’s paired *t*-test; mean ± SEM. Symbols: ▪ males, ● females, ♦ males and females, * *p* < 0.05, ** *p* < 0.01, *** *p* < 0.001. Parts of this figure were created with the help of BioRender.com.

### Selective expression of a pathological *App* allele in cholinergic medial septal neurons generates localized Aβ accumulation and leads to cholinergic neuronal loss

Cre recombinase-dependent AAV transgenes that express the familial *App*^NL-G-F^ mutations were constructed (Figure 2A). The viruses *AAV-DIO-App*^NL-G-F^*-P2A-EGFP* or *AAV-DIO-EGFP* (as a control) were injected into the MS of *ChAT-Cre* male and female mice to generate *MS^ChAT^-App*^NL-G-F^ and *MS^ChAT^-GFP* mice that were then allowed to age for 13 to 14 months (Figure 2A); their phenotypes were then compared in detail with *App*^NL-G-F/NL-G-F^ global *knock-in* mice that were aged at the same time (Figure 2D). As an internal control, to determine whether the phenotypes observed in aged *MS^ChAT^-App*^NL-G-F^ mice were predominantly attributable to the putative loss of MS cholinergic neurons or to the deposition of amyloid, we selectively lesioned MS^ChAT^ cells independently of amyloid expression. *AAV-DIO-Caspase* was injected into the MS of *ChAT-Cre* male and female mice, thereby generating *MS^ΔChAT^* animals, which were subsequently aged until 7-8 months post-surgery, making them around 9-10 and 10-11 months old at the time of sleep and behavioral analysis, respectively.

**Figure 2.**
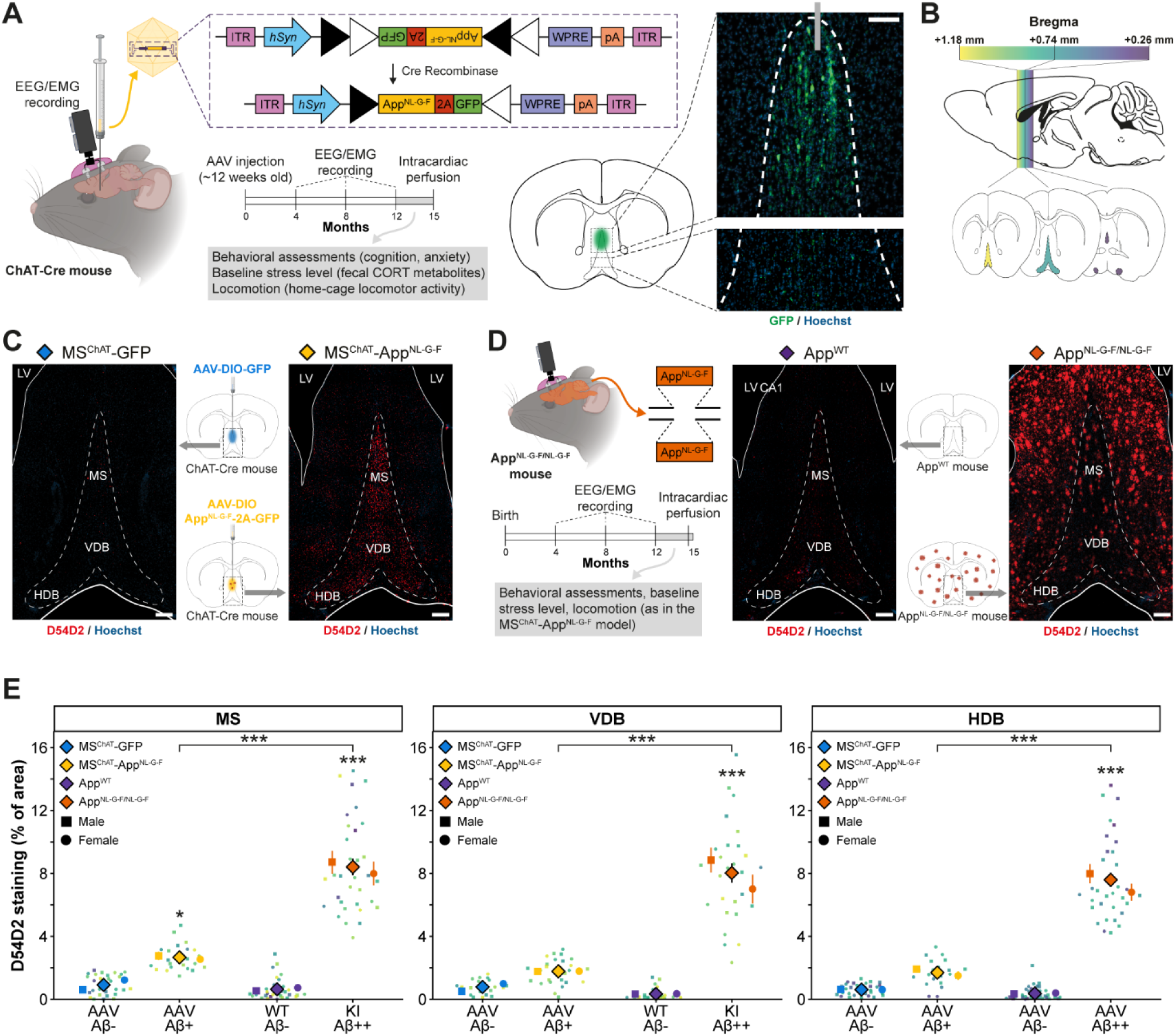
*MS^ChAT^-App*^NL-G-F^ mice exhibit local Aβ deposits, while *App*^NL-G-F/NL-G-F^ *knock-in* mice show marked increase in Aβ deposition. (A) *MS^ChAT^-App*^NL-G-F^ mice were generated by injecting *AAV-DIO-App*^NL-G-F^*-2A-EGFP* into the MS of *ChAT-Cre* mice. EEG/EMG recordings were performed every four months over the year post-injection to monitor sleep-wake cycles. From 12 months onward, behavioral assessments, locomotor activity monitoring, and baseline stress level measurements were conducted prior to brain collection for histological analyses (scale bar: 100 μm). (B) Serial coronal brain sections spanning the rostrocaudal extent of the MS were analyzed. In plots of quantified immunohistochemical BF data, gradient color-coding indicates the rostrocaudal position of each brain slice from anterior to posterior. (C) Coronal brain sections from *MS^ChAT^-GFP* (left) and *MS^ChAT^-App*^NL-G-F^ (right) mice show Aβ immunostaining (D54D2, red) and cell nuclei (Hoechst, blue) (scale bar: 200 μm). (D) *App*^NL-G-F/NL-G-F^ *knock-in* mice, carrying familial Alzheimer’s disease-linked APP mutations, were monitored using the same longitudinal design as *MS^ChAT^-App*^NL-G-F^ mice, with EEG/EMG recordings every four months from birth and behavioral, locomotor, and stress assessments from 12 months onward. Coronal brain sections from *App^WT^* (control; left) and *App*^NL-G-F/NL-G-F^ (right) mice show Aβ immunostaining (D54D2, red) with Hoechst counterstaining (blue) (scale bar: 200 μm). (E) *MS^ChAT^-App*^NL-G-F^ mice exhibited increased Aβ accumulation in the MS, with no significant changes in the VDB or HDB compared with *MS^ChAT^-GFP* controls. In contrast, *App*^NL-G-F/NL-G-F^ mice displayed robust increases in Aβ deposition across the MS, VDB and HDB relative to *App^WT^* controls. Aβ levels were also higher in *App*^NL-G-F/NL-G-F^ mice than in MS^ChAT^-App^NL-G-F^ animals across all BF regions. See panel B for gradient color-coding. Statistical analysis: linear mixed-model ANOVA with Holm-Bonferroni correction for multiple pairwise comparisons where appropriate (*n* = 4 males/group, *n* = 4 females/group); mean ± SEM. Asterisks on *MS^ChAT^-App*^NL-G-F^ and *App*^NL-G-F/NL-G-F^ groups denote significance relative to their corresponding controls. Symbols: ▪ males, ● females, ♦ males and females, * *p* < 0.05, ** *p* < 0.01, *** *p* < 0.001. Parts of this figure were created with the help of BioRender.com.

For 13- to 14-month-old *MS^ChAT^-App*^NL-G-F^ mice, immunohistochemical analyses employing the amyloid-specific antibodies D54D2 and 6E10 (Figures 2C and S1A, respectively) revealed a marked accumulation of Aβ deposits in the MS (respectively *p* = 0.0452 and *p* < 0.001, Figures 2E and S1B). This amyloid deposition extended beyond the MS into adjacent structures of the BF cholinergic system, including the vertical (VDB) and horizontal (HDB) limbs of the diagonal band of Broca. While D54D2 staining showed only non-significant trend in these regions after multiple-comparison correction (Figure 2E), 6E10 staining revealed significant increases in both VDB and HDB (respectively *p* < 0.001 and *p* = 0.0257, Figure S1B). There was also a significant increase in Aβ deposits, assessed using the D54D2 antibody, in the same BF regions of 13- to 14-month-old *App*^NL-G-F/NL-G-F^ mice (*p* < 0.001 for MS, VDB and HDB; Figures 2D-E), albeit with a greater magnitude than in the *MS^ChAT^-App*^NL-G-F^ model (*p* < 0.001, *MS^ChAT^-App*^NL-G-F^ *versus App*^NL-G-F/NL-G-F^ mice).

Notably, in comparison to *MS^ChAT^-GFP* controls, 14-month-old *MS^ChAT^-App*^NL-G-F^ mice had significantly greater Aβ accumulation in regions innervated by MS^ChAT^ cells (Figures 3A-C). This accumulation was particularly prominent in the CA1, CA2 and hilar areas of the hippocampus (*p* = 0.0065, Figures 3A-B), with additional amyloid deposits detected in the thalamus, amygdala, mPFC, primary cortical areas, and the hypothalamus (Figure 3C). However, no increase in Aβ burden was observed in the medial habenula, olfactory bulb, or entorhinal cortex, despite their cholinergic innervation by the MS (see above), nor in the caudate-putamen, where no cholinergic input from the MS was previously detected (see Supplemental information). There was also some colocalization of amyloid and laminin-labeled blood vessels in the hippocampus of *MS^ChAT^-App*^NL-G-F^and, to a much lesser extent, *MS^ChAT^-GFP* mice that was not observed in the BF (Figures S1C-D). This suggests that MS^ChAT^ cell terminals preferentially innervate hippocampal blood vessels.

**Figure 3.**
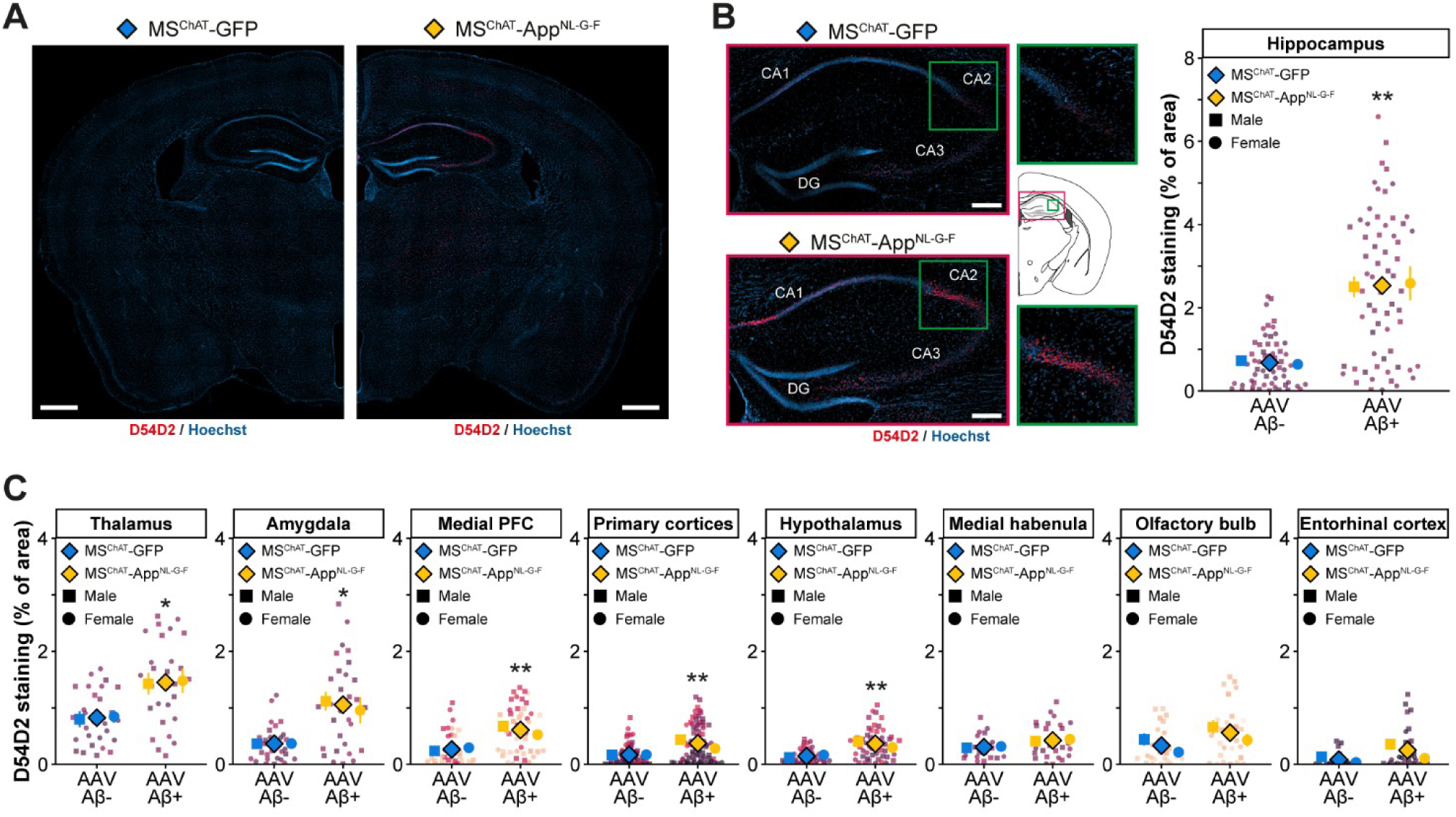
*MS^ChAT^-App*^NL-G-F^ mice exhibit broadcasted Aβ deposits. (A) Coronal whole-brain sections (bregma -1.70mm) showing Aβ immunostaining (D54D2, red) and nuclei (Hoechst, blue) in *MS^ChAT^-GFP* (left) and *MS^ChAT^-App*^NL-G-F^ (right) mice (scale bar: 500 μm). (B) Aβ spread and accumulation in the hippocampus were examined in *MS^ChAT^-App*^NL-G-F^ mice. Coronal hippocampal sections show Aβ immunostaining (D54D2, red) and nuclei (Hoechst, blue) in *MS^ChAT^-GFP* (top) and *MS^ChAT^-App*^NL-G-F^ (bottom) mice (scale bar: 200 μm). *MS^ChAT^-App*^NL-G-F^ mice showed significantly higher Aβ accumulation in the hippocampal regions (CA areas, dentate gyrus, and subiculum) compared with *MS^ChAT^-GFP* mice. See Figure 1C for gradient color-coding. (C) Quantification of Aβ deposition in other projection fields of MS^ChAT^ neurons revealed significantly higher Aβ accumulation in the thalamus, amygdala, medial prefrontal cortex (prelimbic and cingulate cortices), primary sensory cortices (somatosensory, motor, and visual), and hypothalamus in *MS^ChAT^-App*^NL-G-F^ mice compared with *MS^ChAT^-GFP* controls. No significant differences were observed in the medial habenula, olfactory bulb, or entorhinal cortex. See Figure 1C for gradient color-coding. Statistical analysis: linear mixed-model ANOVA with Holm-Bonferroni correction for multiple pairwise comparisons where appropriate (*n* = 3-4 males/group, *n* = 4 females/group); mean ± SEM. Symbols: ▪ males, ● females, ♦ males and females, * *p* < 0.05, ** *p* < 0.01, *** *p* < 0.001.

Both *MS^ChAT^-App*^NL-G-F^ and homozygous *App*^NL-G-F/NL-G-F^ mice exhibited a loss of MS^ChAT^ cells relative to their respective controls, with decreases of 23.1% (*p* < 0.001, Figures 4A, D) and 21.9% (*p* < 0.001, Figures 4B, D), respectively. *App*^NL-G-F/NL-G-F^ mice also had fewer ChAT-expressing neurons in the HDB compared with *App^WT^* controls (Figure 4D). In *MS^ChAT^-App*^NL-G-F^ mice, despite widespread amyloid deposition in the hippocampus, no changes in neuronal (NeuN) staining in hippocampus were observed compared to *MS^ChAT^-GFP* controls (Figure S2).

**Figure 4.**
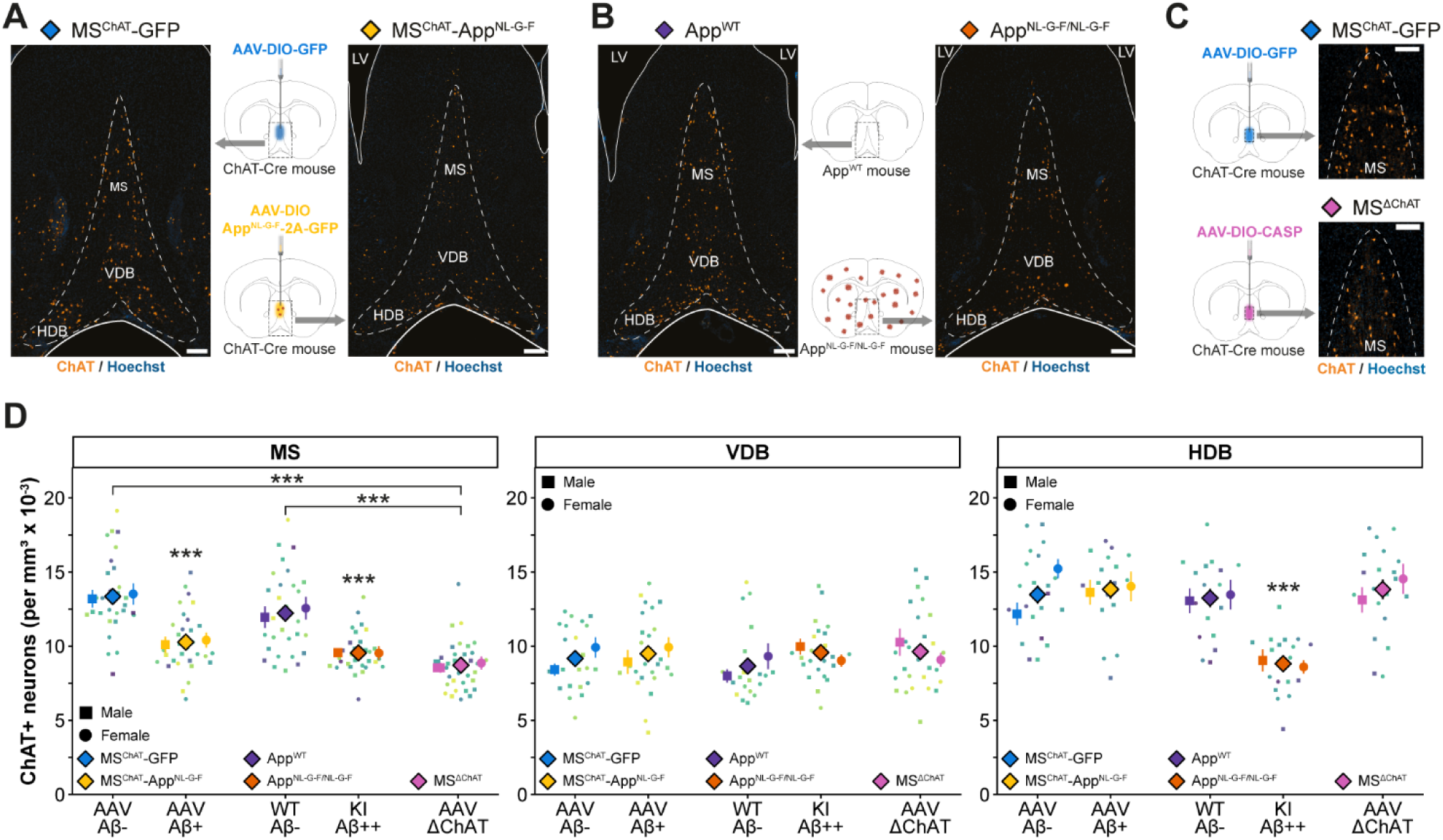
Loss of BF cholinergic neurons in aged *MS^ChAT^-App*^NL-G-F^ and *App*^NL-G-F/NL-G-F^ mice, and in *MS^ΔChAT^* mice. (A-B) Coronal sections showing cholinergic neurons (ChAT, orange) and nuclei (Hoechst, blue) in the MS, VDB, and HDB of (A) *MS^ChAT^-GFP* (control; left) and *MS^ChAT^-App*^NL-G-F^(right) mice and (B) *App^WT^* (control; left) and *App*^NL-G-F/NL-G-F^ (right) mice (scale bar: 200 μm). (C) Coronal MS sections showing cholinergic neurons (ChAT; orange) and nuclei (Hoechst, blue) in *ChAT-Cre* mice after MS-targeted injection of *AAV-DIO-EGFP* (*MS^ChAT^-GFP*, control; top) or *AAV-DIO-CASP* (*MS^ΔChAT^*; bottom) (scale bar: 100 μm). (D) *MS^ChAT^-App*^NL-G-F^ mice exhibited a significant reduction in cholinergic neuron number in the MS compared with *MS^ChAT^-GFP* mice, while the VDB and HDB were unaffected. *App*^NL-G-F/NL-G-F^ mice showed reduced cholinergic neurons in the MS and HDB relative to *App^WT^* controls, with no change in the VDB. *MS^ΔChAT^* mice also displayed a marked MS-restricted loss of cholinergic neurons compared to *MS^ChAT^-GFP* and *App^WT^* mice. See Figure 2B for gradient color-coding. Statistical analysis: linear mixed-model ANOVA with Holm-Bonferroni correction for multiple pairwise comparisons where appropriate (*n* = 4 males/group, *n* = 4 females/group); mean ± SEM. Asterisks on *MS^ChAT^-App*^NL-G-F^ and *App*^NL-G-F/NL-G-F^ groups denote significance relative to their corresponding controls. Symbols: ▪ males, ● females, ♦ males and females, * *p* < 0.05, ** *p* < 0.01, *** *p* < 0.001.

The MS also contains GABAergic and glutamatergic neurons and synapses^65–67^. Compared to controls, *App*^NL-G-F/NL-G-F^ mice had reduced GAD67 and VGLUT2 immunoreactivity (collective cell bodies and synapses) in the MS, VDB, and HDB (Figures S3A-D). No such changes were observed between *MS^ChAT^-App*^NL-G-F^ and *MS^ChAT^-GFP* mice, however (Figures S3A-D). Overall, in the *MS^ChAT^-App*^NL-G-F^ mice, the intrinsic overproduction of amyloid by MS^ChAT^ neurons over about a year caused the degeneration of about a quarter of them, but the resulting amyloid in the MS did not affect the neighboring non-ChAT cells or synapses.

In the *MS^ΔChAT^* mice, complete ablation of MS cholinergic neurons was not achieved by the caspase expression. Instead, there was a partial neuronal loss of approximately one third of cells (31.7%, *p* < 0.001, Figures 4C-D). The number of cholinergic neurons in adjacent BF regions (*i.e.*, VDB and HDB) of *MS^ΔChAT^* mice remained unaltered compared with *MS^ChAT^-GFP* controls (Figure 4D). Although not anticipated, this partial loss of cholinergic neurons turned out to be fortuitous, as the degree of loss was roughly similar to the loss of cholinergic neurons induced by the familial *App*^NL-G-F^ allele, further enhancing the utility of comparing *MS^ChAT^-App*^NL-G-F^ and *MS^ΔChAT^* mice.

### Aβ-induced disruption of the MS-cholinergic system hinders memory and reduces anxiety-like behavior

Home-cage locomotor activity, measured 13- to 14-months following AAV injections, was only elevated in the dark phase in *App*^NL-G-F/NL-G-F^ mice compared with *App^WT^* controls (Figure 5A). Notably, female mice were consistently more active than males across all groups (see Supplemental information). We also assessed baseline stress levels by ELISA-based quantification of fecal corticosterone metabolites (FCM; Figure 5B). *MS^ChAT^-App*^NL-G-F^ mice showed no significant difference in corticosterone concentrations throughout the day compared to *MS^ChAT^-GFP* controls. By contrast, *App knock-in* mice displayed elevated overall baseline FCM levels relative to their *App^WT^* counterparts (Figure 5B). In all types of mice, females consistently exhibited higher baseline levels of FCM than males (see Supplemental information).

**Figure 5.**
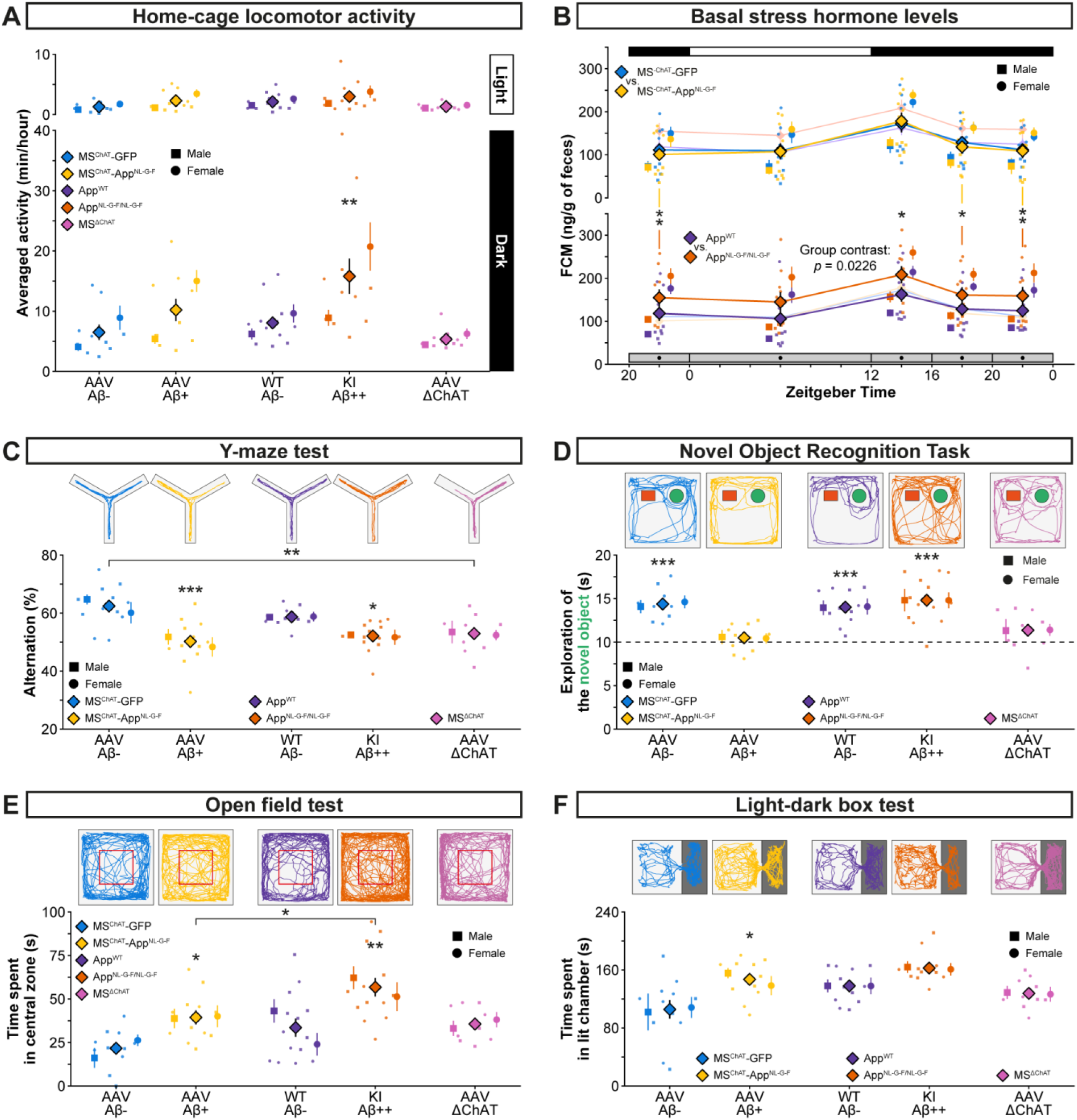
Partially overlapping cognitive and emotional alterations in *MS^ChAT^-App*^NL-G-F^, *App*^NL-G-F/NL-G-F^ and *MS^ΔChAT^* mice. (A) Home-cage locomotor activity monitoring showed that *App*^NL-G-F/NL-G-F^ mice displayed increased locomotor activity relative to *App^WT^* controls during the dark phase. Females were more active than males (see Supplemental information for detailed statistics). (B) Basal stress levels were assessed by quantifying fecal corticosterone metabolites (FCM) across five time windows spanning a 28 h period (gray bar). FCM levels in *MS^ChAT^-App*^NL-G-F^ mice did not differ from *MS^ChAT^-GFP* controls, whereas *App*^NL-G-F/NL-G-F^ mice exhibited significantly elevated levels compared with *App^WT^* counterparts. In both models, female mice showed higher FCM levels than males (see Supplemental information for detailed statistics). (C) Spatial working memory was assessed using the Y-maze test. *MS^ChAT^-App*^NL-G-F^, *App*^NL-G-F/NL-G-F^, and *MS^ΔChAT^* mice, exhibited significantly fewer spontaneous alternations compared to their respective controls, indicating impaired working memory. (D) Non-spatial working memory was evaluated using the novel object recognition task (NORT). During the test phase (maximum 20 s exploration), *MS^ChAT^-App*^NL-G-F^ and *MS^ΔChAT^* mice did not display a significant preference for the novel object, indicating a working memory deficit, whereas *MS^ChAT^-GFP* controls showed a clear preference. In contrast, both *App*^NL-G-F/NL-G-F^ and *App^WT^* mice exhibited significant preference for the novel object. (E) Anxiety-like behavior was assessed using the open field test. *MS^ChAT^-App*^NL-G-F^ mice spent significantly more time in the center compared with *MS^ChAT^-GFP* controls, indicating reduced anxiety-like behavior. *App*^NL-G-F/NL-G-F^ mice showed an even greater increase in center time relative to both *App^WT^* controls and *MS^ChAT^-App*^NL-G-F^ mice, whereas *MS^ΔChAT^* mice did not differ from controls. (F) Anxiety-like behavior was further assessed using the light-dark box test. *MS^ChAT^-App*^NL-G-F^ mice spent significantly more time in the illuminated compartment compared with *MS^ChAT^-GFP* controls, consistent with reduced anxiety-like behavior. In contrast, *App*^NL-G-F/NL-G-F^ and *MS^ΔChAT^* mice did not differ from their respective controls. Statistical analysis: linear mixed-model and standard ANOVAs with Holm-Bonferroni correction for multiple pairwise comparisons where appropriate, and one-sample Student’s *t*-test for NORT (*n* = 5-8 males/group, *n* = 5-7 females/group); mean ± SEM. Asterisks on *MS^ChAT^-App*^NL-G-F^ and *App*^NL-G-F/NL-G-F^ groups denote significance relative to their corresponding controls, except for NORT, where they all denote deviation from chance-level exploration (10 s for each object) Symbols: ▪ males, ● females, ♦ males and females, * *p* < 0.05, ** *p* < 0.01, *** *p* < 0.001.

Behavioral testing was performed 13- to 14-months following AAV injections. Short-term working memory was assessed using the Y-maze paradigm (Figure 5C). Both *MS^ChAT^-App*^NL-G-F^ and *App*^NL-G-F/NL-G-F^ mice performed significantly worse than their corresponding controls *MS^ChAT^-GFP* and *App^WT^* (respectively *p* < 0.001 and *p* = 0.0418), indicating impaired spatial working memory. *MS^ΔChAT^* mice likewise performed significantly worse than *MS^ChAT^-GFP* animals (*p* = 0.0049). In the novel object recognition task (NORT; Figure 5D), *MS^ChAT^-App*^NL-G-F^ mice had no preference for the novel object as opposed to the *MS^ChAT^-GFP* group (*p* < 0.001, *versus* 10 seconds average exploration), suggesting impaired non-spatial object memory; a similar lack of preference was observed in *MS^ΔChAT^* mice. Conversely, *App*^NL-G-F/NL-G-F^ mice, like *App^WT^* controls, explored the novel object more, indicating intact recognition memory (*p* < 0.001, *versus* 10 seconds average exploration). All groups spent comparable amounts of time exploring the two identical objects during the familiarization session of the NORT, confirming equal baseline exploratory behavior (see Supplemental information).

In both the open field and light-dark box tests, *MS^ChAT^-App*^NL-G-F^ mice exhibited reduced anxiety-like behavior compared with controls, with increased time spent in the central zone of the open field (*p* = 0.0444, Figure 5E) and the illuminated chamber of the light-dark box (*p* = 0.0106, Figures 5F). However, in *App*^NL-G-F/NL-G-F^ mice, reduced anxiety-like behavior was observed only in the open field test, compared to both *App^WT^* controls and *MS^ChAT^-App*^NL-G-F^ mice (respectively *p* = 0.0021 and *p* = 0.0444, Figures 5E-F). In contrast, *MS^ΔChAT^* mice did not exhibit a significant reduction in anxiety-like behavior compared to controls.

All animals exhibited broadly comparable exploratory behavior across the conducted behavioral assessments, although group differences were observed in specific measures, including centre entries in the open field (consistent with reduced anxiety-like behavior), and increased distance travelled in the light-dark box for *MS^ΔChAT^* mice compared with *MS^ChAT^-GFP* and *MS^ChAT^-App*^NL-G-F^ mice (see Supplemental information).

### Amyloid pathology restricted to the MS-cholinergic system induces REMS decreases comparable to those in *App*^NL-G-F^ mice

As the mice aged from 4 to 12 months post-AAV injections, we investigated over three time points how amyloid-induced disruption of MS cholinergic neurons affects sleep-wake architecture. In comparison to *MS^ChAT^-GFP* control mice, *MS^ChAT^-App*^NL-G-F^ animals had unchanged amounts of NREMS and WAKE (Figures S4A and S5A). However, *MS^ChAT^-App*^NL-G-F^ mice had fewer (*p* < 0.0042), but prolonged (*p* < 0.0052), NREMS bouts during the light phase, *i.e.* there was enhanced NREMS consolidation (Figures S4B-C). Notably, this same pattern regarding NREMS duration, bout length (*p* = 0.0098), and frequency (*p* = 0.040) was present in homozygous *App*^NL-G-F/NL-G-F^ mice when compared with *App^WT^* controls (Figures S4A-C). Of note, *App*^NL-G-F/NL-G-F^ animals also displayed an overall increase in WAKE time during the light phase (*p* = 0.0175, Figure S5A).

Contrary to the relatively preserved structure of NREMS and WAKE, marked differences in REMS architecture emerged between *MS^ChAT^-GFP* control and *MS^ChAT^-App*^NL-G-F^ mice. Over the year, the amount of REMS in *MS^ChAT^-App*^NL-G-F^ mice lessened relative to baseline controls during the light period, so that by 12 months post-AAV injection *MS^ChAT^-App*^NL-G-F^ mice had clearly less REMS than controls (*p* < 0.001; Figure 6A). During the dark phase, reduced REMS amount was also observed in *MS^ChAT^-App*^NL-G-F^ animals compared to controls (*p* = 0.0256). Moreover, REMS episodes in these mice became strikingly more consolidated: REMS bouts were longer during the light phase (*p* = 0.0278, Figure 6B), but there were fewer of them across both light and dark periods (respectively *p* < 0.001 and *p* = 0.0062, Figure 6C). During a year, presumably as the MS cholinergic neurons were slowly becoming compromised, all these REMS changes (less REMS, longer and fewer bouts) were phenocopied in *App*^NL-G-F/NL-G-F^ mice relative to *App^WT^* controls (Figures 6A-C). Nine- to ten-month-old *MS^ΔChAT^* mice also showed reduced REMS duration and fewer REMS bouts in the light phase compared to *MS^ChAT^-GFP* controls (Figures 6D, F).

**Figure 6.**
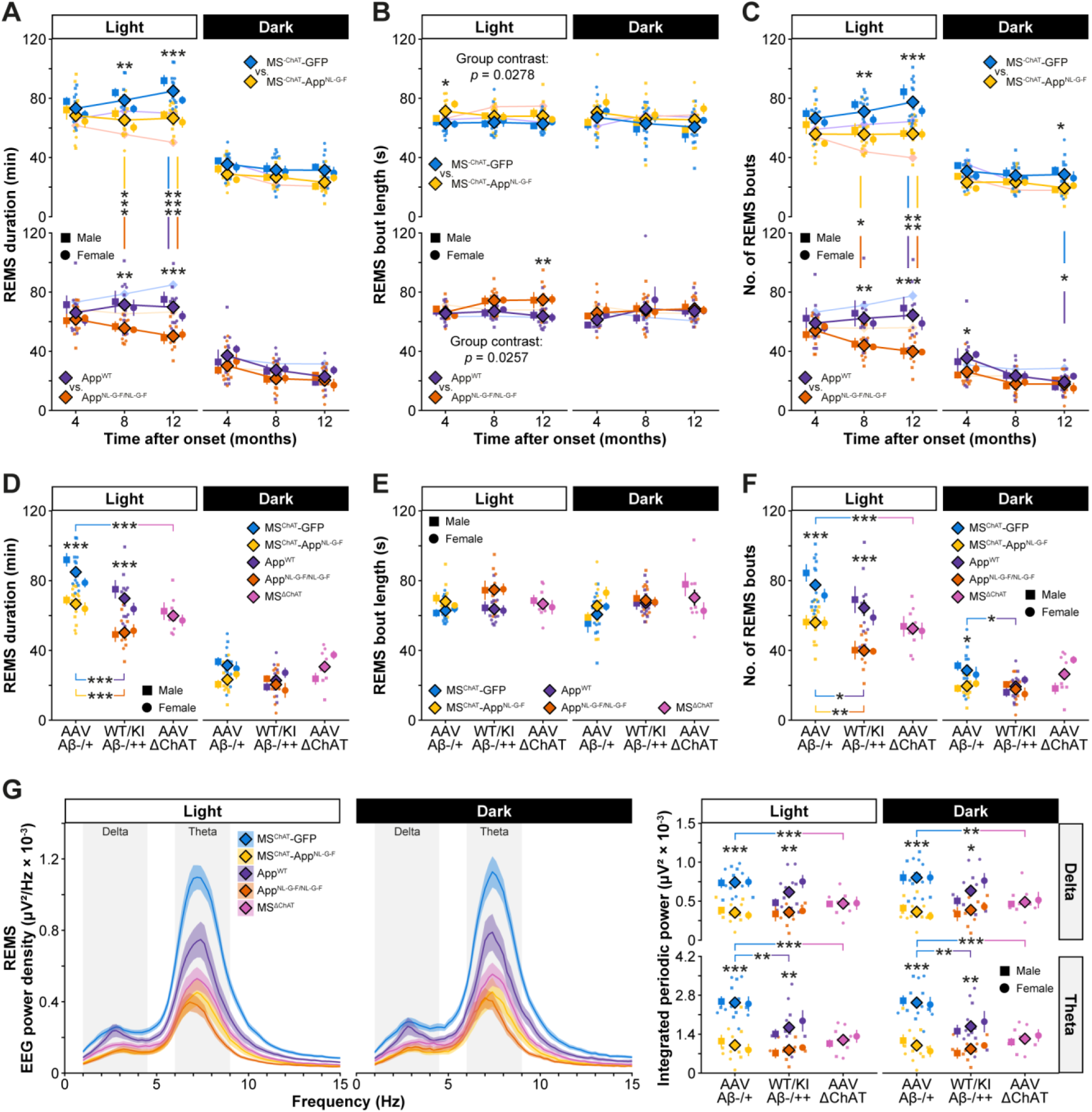
Comparable REMS phenotypes in aging *MS^ChAT^-App*^NL-G-F^ and *App*^NL-G-F/NL-G-F^ mice, and in *MS^ΔChAT^* mice. (A) During the light phase, REMS duration in *MS^ChAT^-App*^NL-G-F^ mice diverged from *MS^ChAT^-GFP* controls at 8 and 12 months following AAV injection, the former group showing overall less REMS. A similar pattern was observed in *App*^NL-G-F/NL-G-F^ mice relative to *App*^WT^ controls at matched stages of amyloid pathology progression. At both stages, *App*^NL-G-F/NL-G-F^ mice exhibited less REMS than *MS^ChAT^-App*^NL-G-F^ mice. Notably, *MS^ChAT^-GFP* and *App*^WT^ controls also differed at 12 months. (B) REMS bout length was overall increased during the light phase in both *MS^ChAT^-App*^NL-G-F^ and *App*^NL-G-F/NL-G-F^ mice compared with their respective controls, irrespective of time scale. Time-specific increases were observed at 4 months post-injection in *MS^ChAT^-App*^NL-G-F^ mice and at later stages of pathology in *App*^NL-G-F/NL-G-F^ mice. (C) In *MS^ChAT^-GFP* mice, the number of REMS bouts increased with time following AAV injection during the light phase, whereas it remained stable in *MS^ChAT^-App*^NL-G-F^ mice, resulting in significant differences at 8 and 12 months. During the dark period, differences between these groups emerged only at 12 months post-injection. In *App*^NL-G-F/NL-G-F^ mice, REMS bout number was reduced compared with *App*^WT^ controls at comparable stages of pathology in the light phase, with an additional reduction observed at earlier stages in the dark phase. In the light phase, REMS bout number was also consistently lower in *App*^NL-G-F/NL-G-F^ mice than in *MS^ChAT^-App*^NL-G-F^ mice at matched stages of amyloid progression. *MS^ChAT^-GFP* and *App^WT^* controls also differed at 12 months in both light and dark periods. (D) Six months after AAV injection, *MS^ΔChAT^* mice displayed reduced REMS duration during the light phase compared with *MS^ChAT^-GFP* mice at 12 months of amyloid pathology. (E) REMS bout length did not differ between *MS^ΔChAT^* mice and other groups. (F) During the light phase, the number of REMS bouts was reduced in *MS^ΔChAT^* mice compared with *MS^ChAT^-GFP* controls. (G) EEG spectral analysis (FOOOF) revealed decreased integrated periodic delta (1-4.4 Hz) and theta (6-9 Hz) power in both light and dark phases in *MS^ChAT^-App*^NL-G-F^ and *App*^NL-G-F/NL-G-F^ mice relative to their respective controls, as well as in *MS^ΔChAT^* mice compared with *MS^ChAT^-GFP* controls. Theta power also differed between *MS^ChAT^-GFP* and *App*^WT^ controls in both phases. Statistical analysis: linear mixed-model and standard ANOVAs with Holm-Bonferroni correction for multiple pairwise comparisons where appropriate (*n* = 5-8 males/group, *n* = 5-8 females/group); mean ± SEM. Symbols: ▪ males, ● females, ♦ males and females, * *p* < 0.05, ** *p* < 0.01, *** *p* < 0.001.

To determine any changes in aperiodic and periodic spectral power of the different vigilance states, we applied a “Fitting Oscillations & One-Over-F” (FOOOF)-based spectral analysis^72,73^. FOOOF analysis of *MS^ChAT^-App*^NL-G-F^ mice at 12 months post-AAV injection revealed significant reductions in delta- and theta-band periodic power (1-4.4 Hz and 6-9 Hz, respectively) during REMS across both phases compared to similarly aged *MS^ChAT^-GFP* controls (*p* < 0.001; Figure 6G), with similar reductions during NREMS (*p* < 0.001; Figure S4G) and wakefulness (*p* ≤ 0.0028; Figure S5G). *App*^NL-G-F/NL-G-F^ mice showed a comparable reduction in delta- and theta-band periodic power during REMS relative to *App^WT^* controls (*p* ≤ 0.0127; Figure 6G); during NREMS, only theta-band power was reduced (*p* ≤ 0.0267; Figure S4G), and during wakefulness, only delta-band power was reduced in the dark phase (*p* = 0.0401, Figure S5G). *MS^ΔChAT^* mice showed a broadly similar pattern, with reductions in delta- and theta-band periodic power during REMS (Figure 6G), NREMS (Figure S4G), and wakefulness (Figure S5G) compared with *MS^ChAT^-GFP* mice, reaching significance in most conditions (*p* ≤ 0.0289).

Regarding the aperiodic component, *MS^ChAT^-App*^NL-G-F^ and *App*^NL-G-F/NL-G-F^ mice both showed a significant reduction in EEG offset during REMS, NREMS, and wakefulness compared to their respective controls across both phases, except for NREMS in *App knock-in* mice (Figures S6A-C). *MS^ΔChAT^* mice showed an offset reduction only during REMS (Figure S6A). No significant differences in the aperiodic exponent were observed in any group or vigilance state (Figures S6A-C).

Together, these findings demonstrate that cholinergic dysfunction of MS cells, produced either by lesioning about a third of them independently of pathological amyloid expression in *MS^ΔChAT^* mice, or by compromising their health and killing about a quarter of them by expressing the familial *App*^NL-G-F^ transgene, produce largely convergent disruptions in sleep architecture and EEG spectral organisation, including the reduction of delta- and theta-band periodic power across vigilance states, and mimicking the changes seen in global *App*^NL-G-F/NL-G-F^ mice. Nevertheless, subtle differences in REMS architecture and aperiodic EEG features between *MS^ChAT^-App*^NL-G-F^ and *MS^ΔChAT^* mice suggest that beyond cholinergic loss, amyloid pathology may engage additional mechanisms (see next section).

### Cognitive and REMS alterations of *MS^ChAT^-App*^NL-G-F^ mice can be primarily attributed to the loss of cholinergic neurons, while MS^ChAT^ amyloid pathology drives additional features beyond cholinergic loss

*MS^ΔChAT^* mice exhibited cognitive and REMS alterations closely resembling those observed in *MS^ChAT^-App*^NL-G-F^ mice, including reduced spontaneous alternations in the Y-maze and a lack of novelty preference in the NORT (Figures 5C-D), together with decreased REMS duration (Figure 6D). In *MS^ΔChAT^* mice, the aperiodic offset was selectively reduced during REMS (Figure S6A), further supporting a specific link between cholinergic dysfunction and REMS regulation. In line with this, Kendall partial correlation analyses across models displaying cholinergic deficits (*MS^ChAT^-App*^NL-G-F^, *App*^NL-G-F/NL-G-F^ and *MS^ΔChAT^*) revealed a positive association between the number of MS cholinergic neurons and both REMS duration and theta-band periodic power during the light phase (Figure 7A).

**Figure 7.**
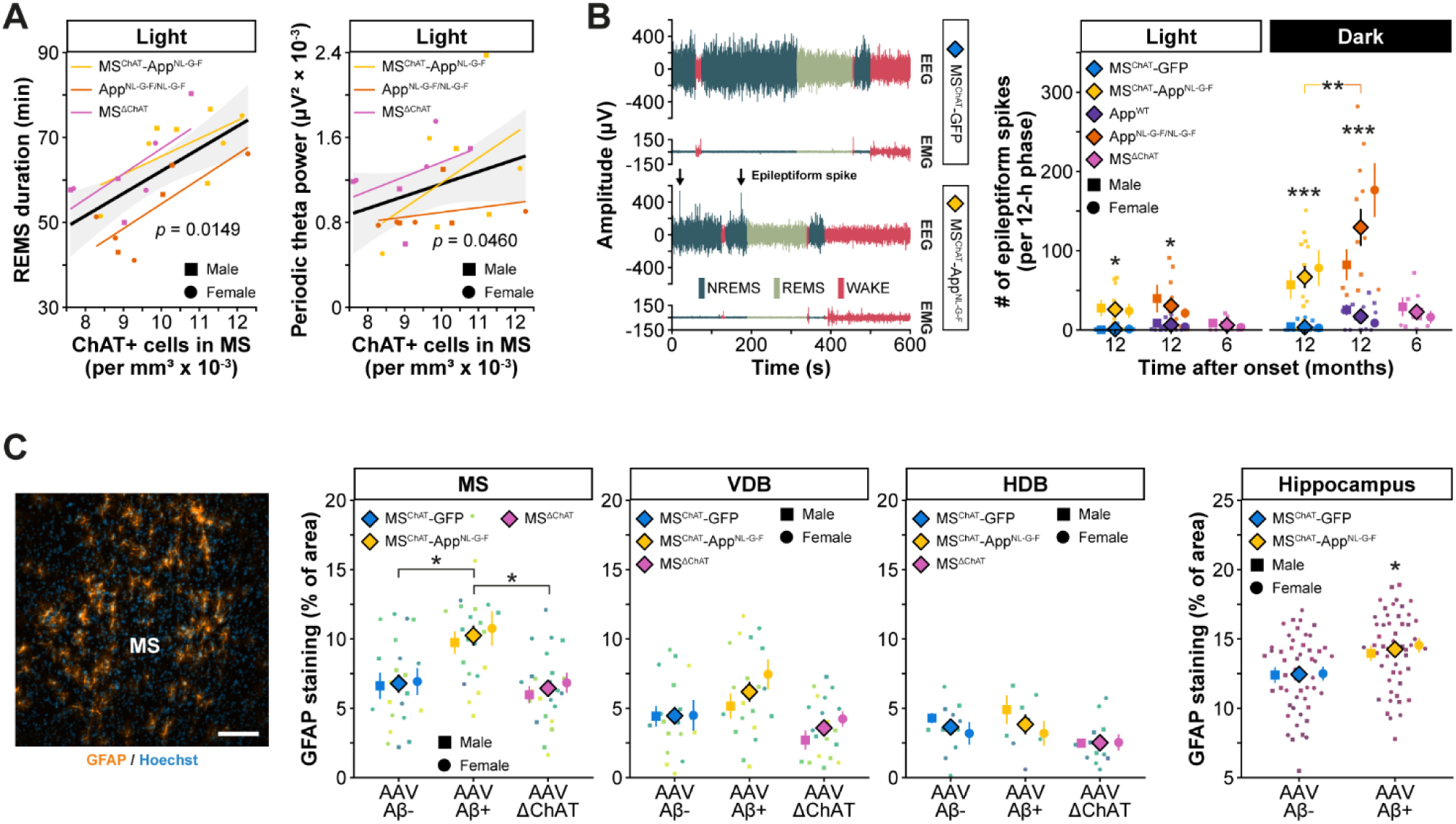
Distinct contributions of amyloid burden and cholinergic loss to phenotypic outcomes. (A) Partial Kendall’s tau correlations across models exhibiting cholinergic loss (*MS^ChAT^-App*^NL-G-F^, *App*^NL-G-F/NL-G-F^, and *MS^ΔChAT^*) revealed positive associations between cholinergic neuron number and both REMS duration and theta power during the light phase. (B) The number of interictal epileptiform spikes was increased in *MS^ChAT^-App*^NL-G-F^ and *App*^NL-G-F/NL-G-F^ mice compared with their respective controls during both light and dark phases. *App*^NL-G-F/NL-G-F^ mice exhibited a higher incidence of spikes than *MS^ChAT^-App*^NL-G-F^ mice, whereas such events were not observed in *MS^ΔChAT^* mice, suggesting a contribution of amyloid pathology to their generation. (C) Coronal sections of the MS showing GFAP immunostaining (astrocytes, orange) and nuclei (Hoechst, blue) (scale bar: 100 μm). *MS^ChAT^-App*^NL-G-F^ mice exhibited increased GFAP expression selectively in the MS compared with *MS^ChAT^-GFP* and *MS^ΔChAT^* mice, consistent with the local accumulation of amyloid, whereas no increase was observed in the VDB or HDB. In the hippocampus, where amyloid is also distributed in *MS^ChAT^-App*^NL-G-F^ mice, GFAP staining was similarly elevated compared with *MS^ChAT^-GFP* controls. Statistical analysis: linear mixed-model and standard ANOVAs with Holm-Bonferroni correction for multiple pairwise comparisons where appropriate, and partial Kendall’s tau correlations controlling for group and sex (*n* = 4-8 males/group, *n* = 4-8 females/group); mean ± SEM. Symbols: ▪ males, ● females, ♦ males and females, * *p* < 0.05, ** *p* < 0.01, *** *p* < 0.001.

We also looked at several amyloid-associated features: epileptiform activity^74–77^, and astrocyte activation^78,79^. Both *MS^ChAT^-App*^NL-G-F^ and *App*^NL-G-F/NL-G-F^ mice had significantly increased epileptiform spike activity compared to their respective controls, approximately 66.77 ± 13.96 and 129.42 ± 23.44 spikes per 12-h dark period, respectively (Figure 7B), whereas *MS^ΔChAT^* mice, which lack amyloid deposition, did not exhibit astrocyte activation or spikes, indicating a specific contribution of amyloid to these network abnormalities.

The amyloid deposition in *MS^ChAT^-App*^NL-G-F^ mice was associated with a localized increase in GFAP immunoreactivity within the MS, which was not observed in adjacent BF regions or in *MS^ΔChAT^* mice, and was also evident in the hippocampus in line with amyloid distribution (Figure 7C).

Together, these findings support a primary role for MS cholinergic neuron loss in producing cognitive and REMS alterations, while highlighting additional, distinct contributions of amyloid pathology originating from MS cholinergic cells to hippocampal network dysfunction and astrocyte activation.

## DISCUSSION

In this study, we show that inducing a focal AD-like pathology in a small population of ACh-producing subcortical neurons leads, as mice age, to some of the typical changes that mark the prodromal phase of AD, such as the gradual loss of REMS. Remarkably, many phenotypic traits of ageing *App*^NL-G-F^ mice, a model for familial AD in which REMS disorder is a known feature, can be reproduced by inducing amyloid pathology solely within MS^ChAT^ cells. ACh released from these neurons into cortical and hippocampal circuits is crucial for sustaining neuronal network function during a wide variety of behaviors^6,7,12^. Our results are threefold: first, a model depicting the gradual decline and selective vulnerability to amyloid of MS^ChAT^ cells as they age and the broadcasting of this amyloid produced within them brain-wide, which could be of relevance in human pathology; second, how loss of ACh from MS cells gradually degrades REMS generation, the theta rhythm, and the regulation of cognition; and third, an independent genetic lesioning of MS^ChAT^ cells, showing that it is primarily the loss of these cells (and thus acetylcholine) that likely drives the aberrant REMS and memory phenotypes, whereas deposited/released amyloid contributes additional and distinct effects, particularly on hippocampal network activity and astrocyte activation.

The *App*^NL-G-F^ allele causes neurodegeneration in knock-in rats and organoids^80,81^. But little cell death has been reported in *App*^NL-G-F^ homozygous mice^71^. We found, however, that a subset (around 22%) of MS^ChAT^ cells degenerate in *App*^NL-G-F^ mice, and that this cell death is reproduced when amyloid is selectively present in MS^ChAT^ neurons as they age. Indeed, human cholinergic BF neurons are unusual in having high amounts of intraneuronal Aβ-accumulation, perhaps as Aβ oligomers, even in young adults^21,62^. Similarly, in the *5XFAD* mouse line, another model which expresses familial APP mutations, amyloid plaques initially form in the “Papez circuit” (mamillary bodies and medial septal neurons) before appearing in other areas^82^. In the *MS^ChAT^-App*^NL-G-F^ mice, we found that the amyloid within the mouse MS^ChAT^ neurons killed about 23% of them during just over a year of expression, yet the local amyloid deposits had no discernable effect on the neighboring glutamate and GABA cells/synapses in the MS. The cell death remained cell-autonomous. Thus, our findings confirm that MS^ChAT^ cells are especially susceptible to amyloid toxicity.

Intriguingly, the amyloid deposits in *MS^ChAT^-App*^NL-G-F^ mice are not only confined to the MS but also extend into the hippocampus (especially in the CA pyramidal cell layers and dentate hilus), and other areas innervated by MS^ChAT^ cells. Whether the amyloid remains inside the axons of MS^ChAT^ neurons or is extracellular, and, if the latter, whether it is released by living MS^ChAT^ cells is unclear. For other types of neurons, amyloid and/or APP can be transported along axons to terminals and released from synaptic vesicles during neuronal activity^83–86^. The anatomical distribution of amyloid in our model, closely matching known septo-hippocampal projections and occurring in the absence of detectable expression in hippocampal neuronal somata, supports a projection-dependent origin of the amyloid rather than amyloid originating from local viral transduction or off-target expression. But despite increased amyloid signal in the hippocampus, NeuN staining did not reveal any significant neuronal loss, indicating that this deposition does not produce overt neurodegeneration in the hippocampus.

The fact that selective ablation of MS^ChAT^ cells alone, through targeted caspase expression, produces cognitive, sleep, and EEG phenotypes largely overlapping with those observed in *MS^ChAT^-App*^NLG-F^ mice raises the possibility that amyloid released both in the BF and from MS^ChAT^ terminals in the hippocampus and other projection areas may not be sufficient *per se* to drive these sleep/behavioral changes. Rather, it is the loss of ACh that seems critical, a notion further supported by Kendall partial correlation analyses across models, showing that the extent of MS cholinergic cell loss is positively associated with REMS duration and theta activity.

Our results support a dichotomy in which early cholinergic dysfunction may be a primary trigger of these phenotypes, while amyloid accumulation, as it progresses and spreads, contributes additional and distinct effects. Consistent with this, while cholinergic loss alone was sufficient to impair both spatial working memory and object recognition, the presence of chronic, widespread amyloid pathology in *App*^NL-G-F/NL-G-F^ mice was associated with a more selective cognitive profile, with preserved recognition memory despite deficits in spatial working memory. Thus, amyloid may differentially shape the expression of cognitive impairments across domains, potentially through long-term network adaptations, while more acute and focal disruption of the MS-hippocampal cholinergic system may be more detrimental to recognition memory.

Features selectively associated with amyloid pathology, epileptiform spiking^74–77^ and astrocyte activation^78,79^, further support the idea that pathology can be broadcast with functional effects. Both *MS^ChAT^-App*^NL-G-F^ and *App*^NL-G-F/NL-G-F^ mice exhibited increased epileptiform spike activity, whereas *MS^ΔChAT^* mice did not. In fact, spiking activity in the *MS^ChAT^-App*^NL-G-^F mice was about half the amount in *App*^NL-G-F/NL-G-F^ mice, suggesting that the broadcasted amyloid has a significant impact on network activity. Amyloid originating from cholinergic septal cells was accompanied by increased GFAP immunoreactivity in the MS and hippocampus, consistent with astrocyte activation linked to amyloid deposition^78,79^. In humans, the released amyloid from MS^ChAT^ terminals could initiate further pathology in other neurons, possibly via prion-like mechanisms^87^, thereby amplifying and diversifying disease manifestations. We also cannot exclude that additional neuromodulatory systems, such as noradrenergic inputs from the locus coeruleus, which also degenerate early in Alzheimer’s disease^88^, may also produce similar phenotypes. However, genetic lesioning of noradrenergic locus coeruleus neurons in mice does not influence gross features of NREMS and REMS or their amounts^89^, but it is expected that loss of noradrenergic LC cells would diminish cognitive abilities^90^.

In the hippocampus, a portion of the amyloid originating from the MS^ChAT^ axons appears to associate with vascular structures, as identified by laminin staining. This suggests that the MS^ChAT^ cells innervate some blood vessels. Indeed, ACh facilitates cerebral vasodilatation and enhances blood flow^91^; on the other hand, reduced vascular reactivity hampers Aβ clearance, potentially contributing to cerebral amyloid angiopathy (CAA)^92^. Supporting this, when cholinergic neurons throughout the forebrain in mice were lesioned with mu-saporin, the loss of cholinergic innervation exacerbated neurovascular impairments and CAA progression in the cortex and hippocampus^93,94^.

Acetylcholine is important for REMS: mice with a double knockout of m1 and m3 muscarinic receptor genes have no REMS^36^. Our study also highlights MS^ChAT^ cells as key drivers of REMS, theta rhythmicity, and cognitive processing. Similar to cholinergic cells in other basal forebrain nuclei^8,95^, we found that MS^ChAT^ cells are specifically active in WAKE and REMS, as also reported by others^96^. This pattern of activity aligns with their involvement in theta oscillations during these states, as evidenced by the diminished theta power density in both WAKE and REMS episodes of *MS^ChAT^-App*^NL-G-F^ and *MS^ΔChAT^* mice, which also experienced significantly less REMS. Although REMS generation is classically thought to be primarily generated in the brainstem^97^, and certainly this contains a core part of the REMS circuitry that generates muscle atonia and degenerates in Parkinson’s disease^97,98^, many other REMS-generating components have now been found throughout the brain, including in neocortex, basal ganglia, hypothalamus, as well as GABA cells in the medial spetum^44,68,99–104^. MS^ChAT^ neurons can now also be added to this list. Importantly, REMS was not abolished but only partially lost in MS^ChAT^-App^NL-G-F^ mice. This is consistent with the distributed REMS-generating model, whereby forebrain cholinergic systems, including MS^ChAT^ neurons, modulate the quantitative expression and electrophysiological features of REMS.

In our study, control *MS^ChAT^-GFP* mice on a C57BL/6 background exhibited an age-related increase in REMS up to 15 months of age, consistent with previous reports showing preserved or increased REMS at middle age in C57BL/6 mice, with declines occurring later in life^105^. The REMS deficit in *MS^ChAT^-App*^NL-G-F^ mice increases with age, presumably reflecting the gradual degeneration of MS^ChAT^ cells. Lesions of the MS abolish hippocampal theta^106^. Theta activity, including during REMS, is thought to be governed by MS^GABA^ cells that project to the hippocampus^66–68^. Opto-inhibiting MS^GABA^ cells reduces hippocampal theta^68^, but opto-stimulation of MS^ChAT^ cells alone failed to elicit hippocampal theta^96^. Despite this, our results indicate that MS^ChAT^ cells contribute to a component of theta rhythm generation.

While other cell-circuit candidates responsible for the reduced REMS phenotype in *App*^NL-G-F^ mice have been proposed, in particular REMS-generating MCH cells in the lateral hypothalamus and ACh neurons in the pontine brainstem^49,107^, these associations were correlative and did not definitively implicate those cell types in mediating the REMS decline. Unexpectedly, our results reveal that MS^ChAT^ neurons may also influence NREMS, accompanied by alterations in EEG spectral power, although the underlying mechanism remains unclear. NREMS disturbances have been observed in *App*^NL-G-F^ (although modest) and in other genetic mouse models^60,108^, highlighting the broader impact of AD-related pathology on sleep-wake regulation. The fact that, however, in all three mouse models examined (*MS^ChAT^-App*^NL-G-F^, *App*^NL-G-F^, and *MS^ΔChAT^*), the remaining REMS bouts were overall longer but less frequent, although the increase in bout length was not consistently observed in *MS^ΔChAT^* mice, may suggest a reorganization within the sleep circuitry, potentially reflecting adaptative changes aimed at preserving aspects of REMS physiology.

Importantly, the question arises about the consequences of less REMS, and also has reduced theta and delta power. Although in mice REMS is dispensable in a laboratory environment^36^, in humans lower REMS percentages are linked to higher all-cause, cardiovascular, and non-cancer mortality^40^. Both REMS and theta rhythms are hypothesized to contribute to enhancing cognition^41–43^, regulating emotions and reducing stress responses^44–46^, processes that are compromised in AD and in individuals living with mild cognitive impairment, as well as in animal models exhibiting widespread Aβ accumulation^47–61^.

In humans, EEG slowing during REMS may represent a neurophysiological marker distinguishing older adults with memory impairment from cognitively intact peers^61,109–113^. However, our findings do not indicate a selective shift in oscillatory slowing *per se*, but rather a widespread reduction in aperiodic offset across REMS, NREMS, and wakefulness in *MS^ChAT^-App*^NL-G-F^ mice, with a similar but more state-dependent pattern in *App*^NL-G-F/NL-G-F^ mice. As the aperiodic offset reflects broadband neural activity and is considered a proxy of global cortical population firing and background cortical gain, these results suggest a reduction in baseline cortical excitability^72^. In line with this interpretation, loss of basal forebrain cholinergic input is known to reduce cortical and hippocampal excitability and population spiking, providing a mechanistic basis for a broadband downward shift in spectral power^15,24^. Notably, *MS^ΔChAT^* mice also exhibited a reduction in aperiodic offset, albeit restricted to REMS, indicating that cholinergic loss alone is sufficient to impact this parameter in a state-dependent manner, whereas the more global reduction observed in *MS^ChAT^-App*^NL-G-F^ mice likely reflects the combined effects of cholinergic dysfunction and amyloid pathology.

Aperiodic components (power decreasing with increasing frequency) vary across vigilance states, supporting a physiological basis for state-dependent modulation of background activity^73^, and are reliably captured using spectral parameterization approaches that dissociate aperiodic and oscillatory components^73^. In the context of amyloid-driven network alterations, both modeling and human MEG/EEG studies indicate the coexistence of regional hyper- and hypoexcitability along the Alzheimer’s disease continuum, with inhibitory or homeostatic mechanisms dominating in some circuits and leading to net reductions in broadband activity that are spatially and state dependent^114^. Thus, rather than reflecting canonical EEG slowing or a uniform state of hyperexcitability, the present results point to a cholinergic-dependent reduction in state-dependent cortical activation, modulated by amyloid pathology, and consistent with emerging views of Alzheimer’s disease as a dynamic condition characterized by region- and stage-specific imbalances in excitation and inhibition^114,115^.

Both *MS^ChAT^-App*^NL-G-F^ and *App*^NL-G-F^ mice exhibited reduced anxiety-like behavior, consistent with previous reports in the *3xTg* transgenic model of AD^116^. Because *MS^ΔChAT^* mice showed no change in anxiety, this effect is likely attributable to amyloid in the ventral hippocampus, a region known to regulate anxiety. In our model, the progressive loss of MS^ChAT^ neurons is expected to markedly diminish cholinergic input to the hippocampus, particularly along the septo-ventral hippocampal axis where we observed dense MS^ChAT^ innervation. Given that septo-ventral hippocampal cholinergic projections are critically involved in REMS-related theta activity, reduced MS cholinergic tone in our model likely underlies the observed REMS alterations, whereas emotional phenotypes may require a more pronounced cholinergic deficit or the additional presence of amyloid pathology.

MS^ChAT^ neurons enhance spatial and non-spatial working memory^117,118^, and their inputs to the dorsal hippocampus seem more strongly implicated in these processes^119^. Inhibition or ablation of MS^ChAT^ projections to the dorsal CA2 region of the hippocampus (where we observed prominent amyloid deposition) impairs social cognition in mice^120,121^, which can be restored through the pharmacological increase of cholinergically-driven theta oscillations^120^. Collectively, these behavioral observations underscore the central role of MS^ChAT^-hippocampal circuit dysfunction in mediating alterations in risk assessment, working memory, and social cognition. These impairments mirror core symptoms seen in AD patients, including impulsivity, diminished threat perception, short-term memory deficits, and socially inappropriate behavior^122^.

Our findings imply that slow disruptions in MS^ChAT^ neurons in the early stages of AD gradually reduce REMS, alter NREMS architecture, decrease EEG theta and delta power (with region- and state-specific effects), and produce cognitive impairments and selective alterations in emotional behavior. As suggested by the specific caspase-induced lesions of MS^ChAT^ cells, neurodegenerative insults affecting these neurons are sufficient to recapitulate key cognitive and sleep phenotypes associated with excessive amyloid production, although emotional alterations appear to be more dependent on the presence of amyloid pathology. Therefore, preserving MS^ChAT^ cells integrity in a variety of neurodegenerative diseases could help slow decline of patients. Encouragingly, choline supplementation to pregnant *App*^NL-G-F^ mice reduces learning and memory impairments and brain Aβ deposition in progeny^123^; and a year-long administration of an acetylcholinesterase inhibitor protected against prodromal atrophy of MS^ChAT^ cells in AD patients^124^. This might suggest that keeping MS^ChAT^ neurons healthier for longer in individuals with AD, or even replacing them, like the approaches being used for dopamine cell loss in Parkinson’s disease, may provide an improved palliative therapy route that could be combined with amyloid-targeting treatments to enhance clinical outcome in AD.

In conclusion, we have provided a circuit-based mechanism and animal model for the prodromal phase of AD, highlighting the selective vulnerability of acetylcholine neurons and offering insight into how early deficits may arise during the prodromal stages of the disease. Our findings support a primary role for a slow loss of MS cholinergic neurons producing cognitive and REMS alterations, while highlighting additional, distinct contributions of amyloid pathology, broadcast from MS cholinergic cells, to hippocampal network dysfunction and astrocyte activation. Our findings underscore the interest in revitalizing the cholinergic hypothesis of AD^5,13,24,25^, and suggest that the cholinergic system remains a salient target for the development of disease-modifying interventions^5^.

## METHODS

### KEY RESOURCES TABLE

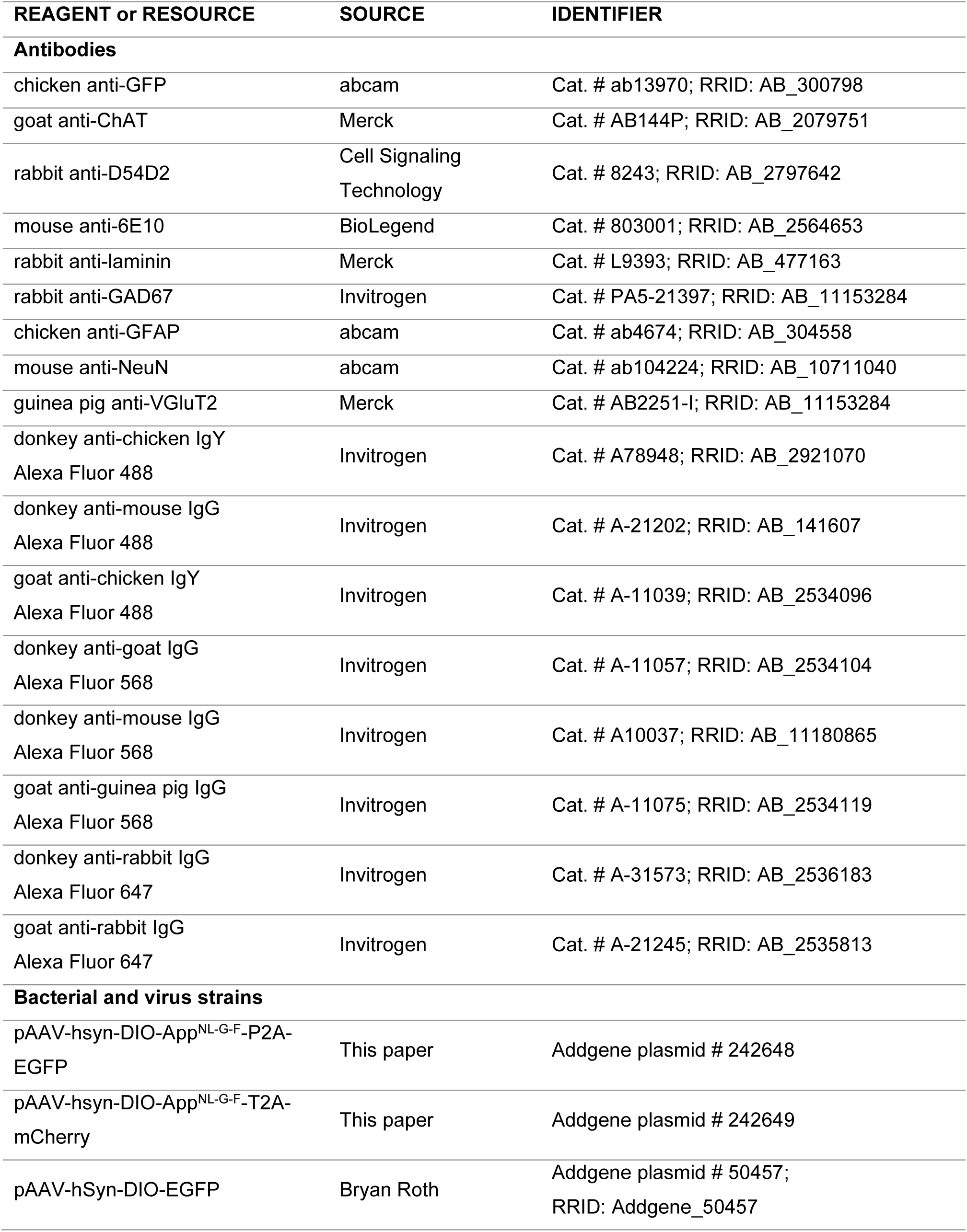
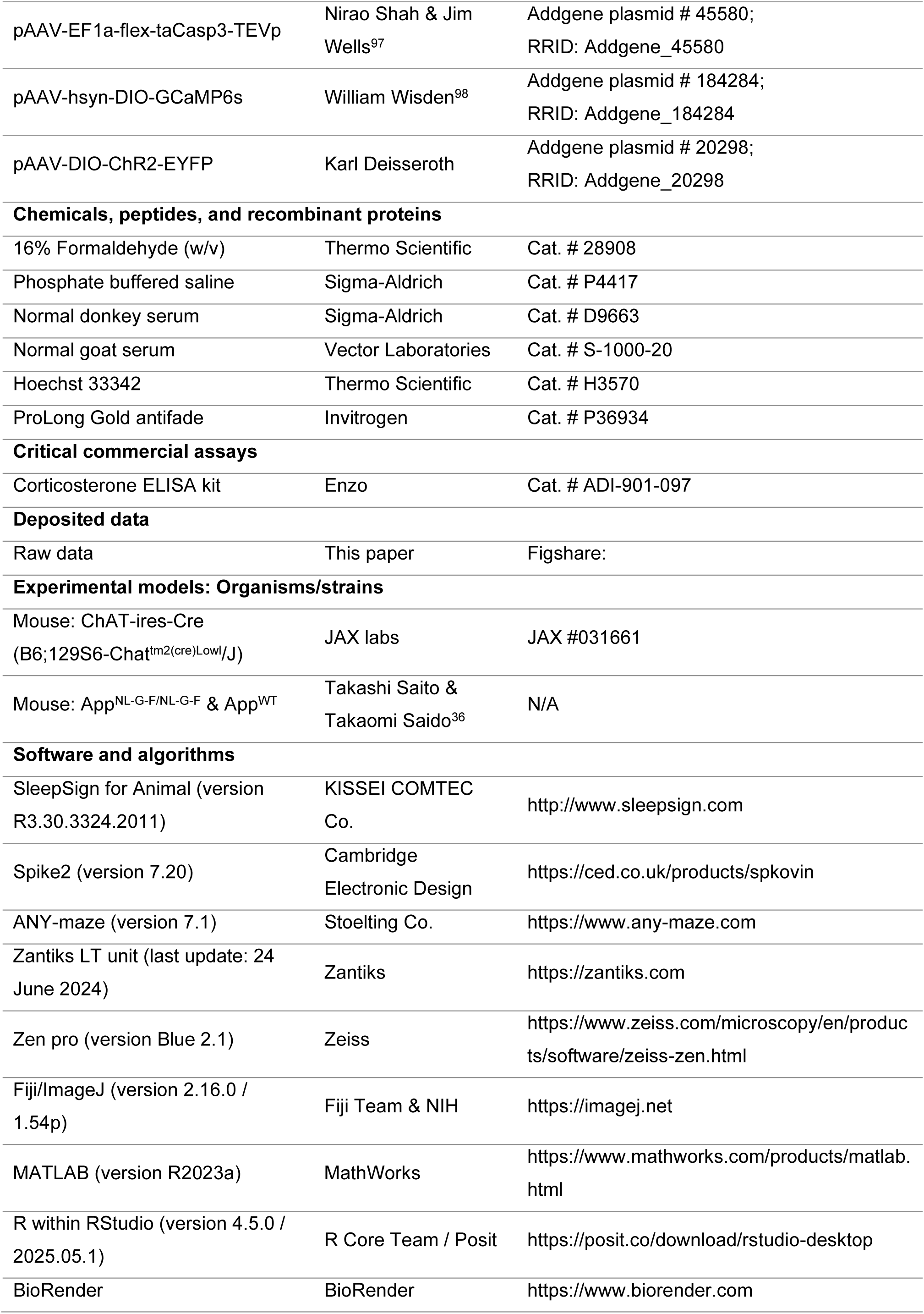
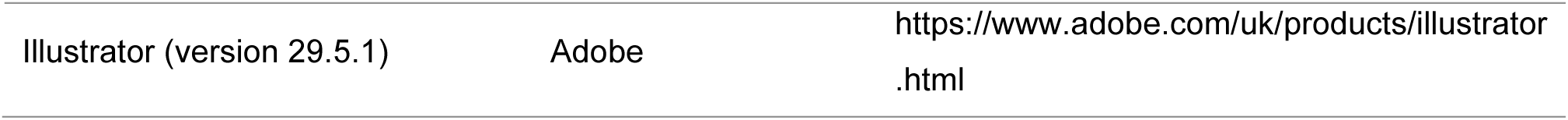

### RESOURCE AVAILABILITY

#### Lead contact

Further information and requests for resources and reagents should be directed to and will be fulfilled by the lead contact, William Wisden.

#### Materials availability

AAVs and plasmids generated in this study have been deposited at Addgene. The accession numbers are listed in the key resources table.

#### Data and code availability

All raw data generated during this study and used to calculate the results reported herein, along with the results of all statistical analyses conducted, as well as bespoke scripts used for sleep analysis, calcium photometry, and histological analyses (developed in MATLAB, R, and Fiji/ImageJ), will be deposited in a public repository upon publication. The accession number will be listed in the key resources table.

### EXPERIMENTAL MODEL DETAILS

We used adult *App*^NL-G-F^ homozygous mice with the Swedish (KM670/671NL), Beyreuther/Iberian (I716F) and Arctic (E22G) *knock-in* mutations in the *APP* gene (these mice were a gift from Takashi Saito, Institute of Brain Science, Nagoya City University Graduate School of Medical Sciences, and Takaomi Saido, RIKEN Centre for Brain Science, Wako, Saitama, Japan)^71^. The *ChAT-ires-Cre* mice (*B6;129S6-Chat^tm^*^2^*^(cre)Lowl^*/J) were from JAX labs (JAX #031661, a gift of Bradford Lowell, Beth Israel Deaconess Medical Center, Harvard)^125^. Animals were maintained under a 12-h light/12-h dark cycle in a controlled environment (21 ± 1 °C; 55 ± 10% relative humidity) with *ad libitum* access to food and water. Environmental enrichment was provided in the form of PVC tunnels and wooden chew blocks. For all mouse strains, males and females were used in all experiments. All procedures were approved by the Animal Welfare Ethical Review Body at Imperial College London. Experiments were performed in accordance with ARRIVE guidelines and the UK Home Office Animal Procedures Act (1986).

### METHOD DETAILS

#### AAV constructs and production

In the *pAAV-hsyn-DIO-App^NL-G-F^-P2A-EGFP and pAAV-hsyn-DIO-App^NL-G-F^-T2A-mCherry* AAV transgenes, the *APP^NL-G-F^*-*P2A-EGFP* or *APP^NL-G-F^*-*T2A-mCherry* reading frames, flanked by heterologous Cre recombination sites, are under the control of the human synapsin promoter. The transgenes were made as follows. An *App*^NL-G-F^ reading frame, flanked by *AscI* and *AgeI* sites, was synthesized by Eurofins (Eurofins Genomics UK Limited) and supplied in a pEX-A258 plasmid. The *App*^NL-G-F^ 2267bp band was gel purified from this plasmid as an *AscI-App^NL-G-F^*-*AgeI* fragment, and ligated into an *AAV-hsyn-DIO-P2A-EGFP* transgene vector doubly-digested with *AscI* and *AgeI*, to generate *AAV-hsyn-DIO-App^NL-G-F^-P2A-EGFP*. A similar procedure generated the *pAAV-hsyn-DIO-App^NL-G-F^-P2A-mCherry* transgene. The transgenes also contained a woodchuck post-transcriptional regulatory element (WPRE) and polyadenylation sequences. These plasmids and their sequences, *pAAV-hsyn-DIO-App^NL-G-F^-P2A-EGFP and pAAV-hsyn-DIO-App^NL-G-F^-P2A-mCherry,* have been deposited at Addgene (Addgene plasmids 242648 and 242649, respectively).

We also used the following *pAAV* transgene plasmids: *pAAV-hSyn-DIO-EGFP* was a gift from Bryan Roth (Addgene plasmid 50457), *pAAV-EF1a-flex-taCasp3-TEVp* was a gift from Nirao Shah & Jim Wells^126^ (Addgene plasmid 45580), *pAAV-DIO-GCaMP6s* transgene was constructed by us and described previously^127^ (Addgene plasmid 184284), *pAAV-DIO-ChR2-EYFP* was a gift from Karl Deisseroth (Addgene plasmid 20298).

The AAVs consisted of a mixed capsid serotype combining AAV1 and AAV2. To generate AAVs, the adenovirus helper plasmid *pFΔ6*, the AAV helper plasmids *pH21* (AAV1) and *pRVI* (AAV2), along with the *pAAV* transgene plasmids, were co-transfected into HEK293 cells. Following this, the resulting AAV particles were purified in-house using heparin columns^128^.

#### Stereotaxic surgery

Prior to surgical procedures, mice were group-housed and allowed at least one week to acclimatize to the animal facility. Surgeries were conducted at 11-13 weeks of age, after which mice were individually housed for the duration of the *in vivo* experiments. Mice were randomly assigned to either experimental or control groups. Anesthesia was induced and maintained with 2% isoflurane in oxygen, delivered via inhalation. Preoperative analgesia and anti-inflammatory treatment were administered subcutaneously using buprenorphine (0.1 mg·kg^-1^) and carprofen (5 mg·kg^-1^). Animals were then positioned and secured in a stereotaxic apparatus (Angle Two; Leica Microsystems) for the duration of the surgical procedure. Postoperatively, carprofen was provided in the drinking water (5 mg·kg^-1^ per day), and mice were allowed to recover in their home cages for a minimum of 3 weeks prior to further experimentation.

##### AAV Injections

Viral vectors were delivered using a stainless steel 33-gauge internal cannula (13 mm, PST3; Hamilton Company) connected to a 10-μl Hamilton microsyringe. Injections were performed at a rate of 0.1 μl·min^-1^, with the following constructs and volumes: *pAAV-hsyn-DIO-App^NL-G-F^-P2A-EGFP* (1 μl), *pAAV-hSyn-DIO-EGFP* (1 μl), a combination of *pAAV-EF1a-flex-taCasp3-TEVp* and *pAAV-hSyn-DIO-EGFP* (0.5 μl each), *pAAV-DIO-GCaMP6s* (0.5 μl), and *pAAV-DIO-ChR2-EYFP* (1 μl). The cannula was slowly lowered to the target site in the MS (coordinates: 0.05 mm ML, 0.8 mm AP, -4.4 mm DV relative to bregma). Following injection, the cannula was left in place for 5 min before being withdrawn slowly to minimize backflow.

##### Fiber photometry implantation

In mice receiving *AAV-DIO-GCaMP6* injections, a 200-μm diameter, 0.5 NA (numerical aperture) mono-fiber optic cannula (RWD Life Science) was surgically implanted 100 μm above the injection site in the MS and secured with a cured acrylic resin (Simplex Rapid powder and Rapid Liquid, Kemdent). Mice were allowed to recover for 3 weeks prior to photometry recording.

##### Headstage implantation for EEG/EMG recordings

All mice were implanted with two miniature screw electrodes for EEG recording. Electrodes were prepared by soldering ultra flexible PVC lead EEG wires (NUF30-4046BLACK, Cooner Wire) to the screws and positioned at standard locations: frontal cortex (-1.7 mm ML, +1.5 mm AP relative to bregma) and parietal cortex (-1.7 mm ML, +1.0 mm AP relative to lambda). EMG electrodes (stranded stainless-steel wire; AS634, Cooner Wire) were inserted bilaterally into the neck musculature. All wires were soldered to a 7-pin header, which was secured to the skull using a cured acrylic resin (Simplex Rapid powder and Rapid Liquid, Kemdent).

#### Sleep-wake analysis

Continuous EEG and EMG recordings were acquired for a minimum of 48 h using untethered Neurologger 2A devices^129^, which were connected to a 7-pin headstage implant. Data were recorded at a sampling rate of 200 Hz with fourfold oversampling. In MATLAB (MathWorks), EEG signals were processed using a high-pass digital filter (highpass function; cutoff frequency: 0.5 Hz, -3 dB), while EMG signals were filtered using a band-pass digital filter (bandpass function; 5-45 Hz, -3 dB).

##### Sleep scoring

For sleep state classification, 24-h EEG/EMG recordings were analyzed, comprising 12-h light and 12-h dark phases. These recordings were collected 3 times at 4-month intervals following AAV injection or from birth in both *App*^NL-G-F/NL-G-F^ and *App^WT^* mice. For *MS^ΔChAT^* mice, recordings were collected 6 months post-injection. Vigilance states, *i.e.* non-rapid eye movement sleep (NREMS), rapid eye movement sleep (REMS) and wakefulness, were manually scored in 5-s epochs using SleepSign for Animal software (KISSEI COMTEC Co.) by an experienced experimenter blinded to treatment conditions.

##### EEG power spectral analysis

EEG power spectral densities were computed for successive 5-s epochs using the Welch method in MATLAB (pwelch function; frequency range: 1-15 Hz; resolution: 0.2 Hz; non-overlapping Hanning window), a more robust and less noisy variant of fast Fourier transform. Epochs containing artifacts were excluded from spectral analyses. Power spectra were calculated separately for NREMS, REMS, and wakefulness during both light and dark phases. To further characterize spectral features, power spectra were parameterized using the Fitting Oscillations and One-Over-F (FOOOF) algorithm, implemented in Python^73^. This approach decomposes the power spectrum into an aperiodic (1/f-like) component and superimposed periodic oscillatory peaks, allowing separation of broadband background activity from true oscillatory signals. The aperiodic component was described by its offset, reflecting the overall broadband power level, and its exponent, reflecting the slope of the spectrum and the relative distribution of power across frequencies. Periodic (oscillatory) activity was quantified after removal of the aperiodic component by integrating power within predefined frequency bands. Delta (1-4.4 Hz) and theta (6-9 Hz) periodic power were extracted for each vigilance state and phase. In parallel, a conventional spectral analysis was performed. For each epoch, spectral power within each 0.2 Hz EEG frequency bin was normalized to the total spectral power of that epoch and expressed as a percentage of total EEG power. Delta power and theta band power were then calculated by summing normalized power within the respective frequency ranges.

Interictal epileptiform spikes were identified using an automated detection algorithm based on a previously validated method^72^. Briefly, the approach combines band-pass filtering with envelope-based feature extraction to detect transient events meeting amplitude and temporal criteria consistent with epileptiform discharges.

#### Calcium photometry

To assess the spontaneous activity of MS cholinergic neurons across sleep-wake states (NREMS, REMS, and wakefulness), we employed fiber photometry using the genetically encoded calcium indicator GCaMP6s, as previously described^127,130^. Simultaneous EEG/EMG recordings were acquired to accurately determine vigilance states during photometry sessions. Recordings were conducted minimum 3 weeks post-surgery during the light (rest) phase between zeitgeber time (ZT) 0 and ZT6, with each session lasting 3-6 h. Each animal underwent 3 to 4 recording sessions.

##### Photometry recording

A 473-nm diode-pumped solid-state blue laser (ADR-800A, Shanghai Laser & Optics Century Co.) was used to excite GCaMP6s. The excitation light was routed through a single-source fluorescence cube (FMC_GFP_FC, Doric Lenses) *via* an optical fiber patch cord (Ø 200 μm, 0.22 NA, Doric Lenses). Emitted fluorescence passed through the same cube and was transmitted via a multimodal optical patch cord (Ø 200 μm, 0.37 NA, Doric Lenses) connected to the ceramic-implanted fiber optic cannula (Ø 200 μm, 0.37 NA, Doric Lenses) using a split mating sleeve ferrule (Thorlabs). The GCaMP6s emission signal was filtered between 500-550 nm and detected by an amplified photodiode (APD-FC, Doric Lenses). The photodiode output was relayed to a lock-in amplifier (SR810, Stanford Research Systems), which also modulated the laser at 125 Hz. Laser power was maintained at 80 μW at the fiber tip. The demodulated signal was digitized using a CED 1401 Micro Box (Cambridge Electronic Design) and recorded at a sampling rate of 1 kHz using Spike2 software (Cambridge Electronic Design).

##### Neuronal activity analysis

All photometry data were analyzed using MATLAB. Signals were temporally aligned with the EEG/EMG recordings. Analyses focused on 100-s windows enclosing transitions between vigilance states (NREMS to REMS, NREMS to wakefulness and wakefulness to NREMS). The raw fluorescence signal (F) was normalized to a baseline using the formula^127^:

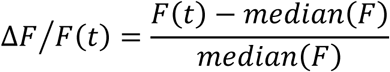

Following ΔF/F normalization, photometry traces were smoothed on a per-transition basis using a Savitzky-Golay filter (3^rd^-order polynomial, 1 s window) implemented in the signal package in R, to reduce high-frequency noise while preserving the slower dynamics of cholinergic signals, and were then averaged across transitions. In some recordings, an initial transient decay in signal intensity was observed; therefore, only sessions with stable photometry signals following this decay period were included in the final analysis.

#### Histological analysis

##### Immunohistochemistry

Mice were transcardially perfused with 4% paraformaldehyde (Thermo Scientific) in phosphate-buffered saline (PBS; Sigma). *MS^ChAT^-App*^NL-G-F^ and *MS^ChAT^-GFP* mice were perfused 13-14 months after AAV injection, whereas *App*^NL-G-F/NL-G-F^ and *App^WT^* mice were perfused at 13-14 months of age, corresponding to comparable durations of amyloid pathology. At this late time point, in *MS^ChAT^-App*^NL-G-F^ mice injected with *AAV-DIO-App*^NL-G-F^*-2A-EGFP*, only sparse and morphologically compromised GFP signal was detectable by immunostaining at the injection site, consistent with the progressive loss of AAV-transduced *ChAT-Cre* neurons driven by sustained intracellular amyloid burden. *MS^ΔChAT^* mice were perfused 7-8 months post-injection, and *ChAT-Cre* mice used for cholinergic projection mapping were perfused 1 month after AAV injection. Brains were extracted, post-fixed in 30% sucrose in PBS for cryoprotection, and later coronally sectioned (40 μm thickness) using a HM 450 sliding microtome (Thermo Scientific). Free-floating sections were washed 3 times in PBS (5 min each), followed by epitope retrieval in 0.05% Tween-20 in 10 mM sodium citrate buffer (pH 6.0) at 80-85 °C for 30 min. Sections were then cooled at room temperature for 15 min and washed 3 times in PBS (10 min each). Non-specific binding was blocked by incubating the sections for 1 h in PBS containing 0.2% Triton X-100 and 20% normal donkey or goat serum. Sections were incubated overnight at 4 °C on a shaker with one or two primary antibodies diluted in PBS supplemented with 2% normal serum. The following day, sections were washed 3 times in PBS (10 min each), incubated for 1.5 h at room temperature with the appropriate secondary antibody diluted in PBS with 2% normal serum, and washed again 3 times (10 min each). When two primary antibodies were used, each corresponding secondary antibody incubation was performed sequentially, with intermediate PBS washes (3 times, 10 min each), to prevent cross-reactivity. For triple labeling, a third primary antibody was applied, followed on the subsequent day by the corresponding secondary antibody as described. Nuclear staining was performed by incubating the sections in Hoechst (1:5,000; Invitrogen) for 3 min at room temperature, followed by three washes in PBS (5 min each). The primary antibodies used included chicken anti-GFP (1:1,000; abcam, ab13970), goat anti-ChAT (1:200; Merck, AB144P), rabbit anti-D54D2 (1:500; Cell Signaling Technology, 8243), mouse anti-6E10 (1:1,000; BioLegend, 803001), rabbit anti-laminin (1:350; Merck, L9393), rabbit anti-GAD67 (1:500; Invitrogen, PA5-21397), guinea pig anti-VGluT2 (1:200; Merck, AB2251-I), chicken anti-GFAP (1:1000; abcam, ab4674) and mouse anti-NeuN (1:1000; abcam, ab104224). Secondary antibodies (1:500; Invitrogen) included Alexa Fluor 488-conjugated donkey anti-chicken (A78948), donkey anti-mouse (A-21202), and goat anti-chicken (A-11039), Alexa Fluor 568-conjugated donkey anti-goat (A-11057), donkey anti-mouse (A10037), and goat anti-guinea pig (A-11075), as well as Alexa Fluor 647-conjugated donkey anti-rabbit (A-31573) and goat anti-rabbit (A-21245). Sections were mounted on glass slides using ProLong Gold antifade mounting medium (Invitrogen) and cover-slipped.

##### Data analysis and quantification

Images were acquired using a Zeiss AxioObserver inverted widefield microscope, controlled with Zen Pro software (Zeiss) at the Facility for Imaging by Light Microscopy, Imperial College London. Illumination was provided by a Lumencor SpectraX LED light source with the following excitation-emission filters: 390/40-450/40 (Hoescht), 470/40-525/50 (Alexa Fluor 488), 550/25-605/70 (Alexa Fluor 568), and 640/30-690/50 (Alexa Fluor 647). Emission was collected with a sCMOS Hamamatsu ORCA-Flash4.0 camera coupled to objectives selected according to the experiment, including 10x/0.3 Ph1 EC Plan-Neofluar, 20x/0.8 DIC Plan Apochromat, and 40x/0.75 Ph2 EC Plan-Neofluar. For large region acquisitions, tiles were configured to overlap by 10% and manual focus maps were generated. Post-acquisition, images were stitched using Zen Pro software. Histological analyses were conducted using the Fiji distribution of ImageJ. Core preinstalled functions were employed for unbiased quantification of neuronal counts and stained areas, utilizing automatic thresholding algorithms included in the Fiji package. Co-localization analyses were performed using the BIOP JaCoP plugin, allowing for quantitative assessment of signal overlap between markers.

#### Behavioral and baseline stress analysis

All behavioral assessments were conducted 13-14 months post-surgery or afterbirth, and during the animals’ active (dark) phase under red light, specifically between Zeitgeber Time (ZT) 16 and ZT22. Animal behavior was recorded using an overhead camera and analyzed automatically in a genotype-blind manner with the ANY-maze software (Stoelting Co.), except for the light-dark box test which was conducted using the Zantiks LT automated apparatus (Zantiks), ensuring objective and unbiased assessment. Video tracking enabled precise quantification of locomotion, orientation, activity, and total distance traveled.

##### Y-maze

The Y-maze task was employed to evaluate spontaneous alternation behavior, an index of spatial working memory^131^. Each mouse was placed at the center of a Y-shaped maze (arm length: 30 cm, width: 6 cm, wall height: 15 cm), with the orientation of entry arms counterbalanced across subjects. Arm entries and total distance traveled were recorded over a 10-min session. A spontaneous alternation was defined as consecutive entries into three different arms. The percentage of spontaneous alternation was calculated as:

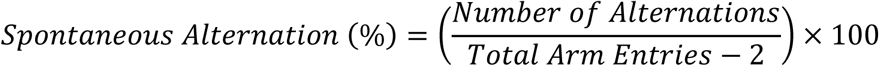

##### Novel object recognition task (NORT)

To assess non-spatial working memory, the NORT was performed as described previously^132^. During the habituation phase, mice were allowed to explore an empty square arena (45 cm × 45 cm, height: 40 cm) for 10 min. This session also served as the open field test for evaluating anxiety-like behavior (see below). Twenty-four hours later, in the familiarization phase, two identical objects were placed 5 cm from the arena walls, and mice were allowed to explore until a total of 20 s of object interaction was reached. This criterion ensured consistent exploration across animals, irrespective of their inherent activity levels. Another 24 hours later, one familiar object was replaced with a novel one, and mice were again allowed to explore both objects until 20 s of cumulative exploration time was reached. Object type and positioning were randomized between animals and experimental groups, and the time to reach the 20 s exploration criterion was recorded.

##### Open field

Anxiety-like behavior was evaluated using the open field paradigm. Mice were placed in the center of a square arena (45 cm × 45 cm, height: 40 cm) and allowed to explore freely for 10 min. The time spent in the central zone, defined as 25% of the total area and considered anxiogenic, were measured, as well as the number of entries into the central zone and the total distance traveled. This session also constituted the initial habituation phase for the NORT (see above).

##### Light-dark box

Bright-light-induced anxiety was assessed using the light-dark box test. The setup included a white bright compartment (23.5 cm × 27 cm, height: 30 cm; ∼150 lux illumination from a translucent floor) connected to a dark compartment (11.5 cm × 27 cm, height: 30 cm) via a 5 cm semi-circle opening. The dark compartment was covered with a removable black semi-translucent lid. Mice were introduced into the center of the dark compartment, facing away from the connecting opening, and allowed to explore both compartments for 5 min. Time spent in the illuminated compartment, the number of transitions between compartments and the total distance traveled were recorded.

##### Home-cage locomotor activity

Spontaneous locomotor activity in the home cage was continuously monitored for an average of 6.8 consecutive days using custom-built devices modeled after a previously published protocol^133^. Each unit consisted of a passive infrared sensor connected to a customized microcontroller (M0 AdaLogger, Adafruit #2796), housed within a 3D-printed casing containing a lithium-ion battery for wireless operation. Devices were placed in the cage on the wire lid, allowing uninterrupted monitoring of activity (total activity duration per min) without disturbing the animals.

##### Assessment of stress hormone levels

Basal stress levels were assessed via fecal corticosterone metabolite (FCM) concentrations using a commercial ELISA kit (ADI-901-097, Enzo), following a previously described protocol optimized and validated in our laboratory^134^. Feces were collected over a 28-h period, divided into five ZT-defined windows: ZT20-ZT0 (final 4 h of dark phase), ZT0-ZT12 (entire light phase), ZT12-ZT16, ZT16-ZT20, and ZT20-ZT0. One week prior to sample collection, animals were habituated to alternating between two uncleaned cages (containing their own scent, enrichment and bedding material), minimizing environmental stress during sampling. At the beginning of the collection period, mice were transferred to the other previously occupied but feces-free cage, and subsequent swaps were performed at each ZT interval. Feces were collected from the previously used cage after each swap and stored at -20 °C until processing. Each sample was homogenized in 5 mL of 95% ethanol per gram of feces, thoroughly ground, and incubated overnight at room temperature. Samples were centrifuged at 3,000 × g for 20 min; the supernatant was collected and centrifuged again at 10,000 × g for 15 min. A 200 μL aliquot of the final supernatant was evaporated to dryness in an open tube under a fume hood. The residue was then reconstituted in 250 μL of assay buffer (provided with the ELISA kit), vortexed, incubated overnight at 4 °C, and stored at -20 °C until analysis. All samples were diluted 1:10 prior to assay and processed in accordance with the manufacturer’s instructions.

### QUANTIFICATION AND STATISTICAL ANALYSIS

All statistical analyses were conducted using R (R Core Team) in RStudio (Posit). *A priori* power analyses were performed prior to the start of the study using previously collected in-lab data to estimate the required sample size for achieving adequate statistical power (typically α = 0.05 and power ≥ 80%). All datasets satisfied the assumptions necessary for parametric testing, including normality and homoscedasticity, assessed through residual inspection and/or standard tests (e.g., Shapiro-Wilk for normality and Levene’s test for equality of variances, where appropriate).With the exception of photometry recordings and the NORT, which were analyzed using paired and one-sample Student’s *t*-tests respectively, all other data were analyzed using either standard ANOVA (via the lm function in base R) when no random effects were present, or linear mixed-effects models (via the lmer function from the lme4 R package) when accounting for random factors, such as repeated measures from the same subject or anatomical variability across brain section locations in histological analyses. In this framework, random intercepts were included for both individual animals and atlas-defined brain section level to account for repeated measurements and residual variability across rostro-caudal sampling positions. Although not all animals contributed data to every section level, this approach allowed us to retain spatially resolved measurements without averaging across sections, while accounting for non-independence of observations.

Fixed effects included experimental group, sex, and time (for repeated measures), modeled as interacting categorical variables, thereby allowing the assessment of group × sex interactions, as well as brain region in the analysis of 6E10/laminin colocalization staining. Given that the primary aim of this study was to define group-related phenotypes across the full cohort, results are presented as pooled male and female comparisons for clarity and interpretability. Full statistical outputs, including sex-stratified analyses and interaction effects, are reported in the Supplementary Information. *Post hoc* pairwise comparisons were conducted using Holm-Bonferroni correction for multiple testing, implemented through the lsmeans and contrast functions from the emmeans R package.

Bivariate associations were assessed using partial Kendall’s tau correlations, controlling for group and sex, as implemented in the R package ppcor.All statistical tests were two-sided, and exact p-values are reported unless otherwise specified. When models included time as a factor, statistical reporting in the main text was restricted to overall group effects to maintain clarity, with detailed pairwise comparisons across time points available in the figures and Supplemental information. No data points were excluded unless predefined technical quality criteria were not met (e.g., insufficient signal quality or failed recordings), in which case the affected measurements were omitted from the corresponding analyses. Sample sizes (n) refer to individual animals unless otherwise indicated. Data are presented as mean ± standard error of the mean (SEM). Full statistical results are provided in the Supplemental information.

## Acknowledgements

Preparation for this article was supported by the UK Dementia Research Institute (award number UK DRI-5004) through UK DRI Ltd, principally funded by the Medical Research Council (W.W., N.P.F.); the Wellcome Trust (220759/Z/20/Z, to N.P.F. and W.W.); and the Medical Research Council (S.W., N.P.F and W.W., APP9329). The Facility for Imaging by Light Microscopy (FILM) at Imperial College London is part-supported by funding from the Wellcome Trust (104931/Z/14/Z) and BBSRC (BB/L015129/1). For the purposes of Open Access, the authors have applied a Creative Commons Attribution (CC BY) license to any Author Accepted Manuscript Version arising.

## Author contributions

M.N. performed behavioral experiments, surgeries and AAV injections, EEG and behavioral analysis, histology, all general data analysis and assembled and designed all figures.

W.B. performed surgeries and calcium photometry analysis.

B.A.S. performed surgeries and helped with histology experiments.

C.Y. helped with histology experiments.

L.L. helped with stress hormone level analysis.

K.J. helped with histology experiments.

S.W. helped with histology experiments.

A.L.V. provided the Neurologger 2A devices.

R.Y. performed all molecular biology and AAV transgene construction, AAV production, mouse genotyping.

M.N., N.P.F and W.W. conceived and designed the experiments. N.P.F. and W.W. co-supervised the project. M.N., N.P.F. and W.W. co-wrote the paper.

## Competing interests

The authors declare no competing interests.

## Supplemental Figures and legends

**Figure S1.**
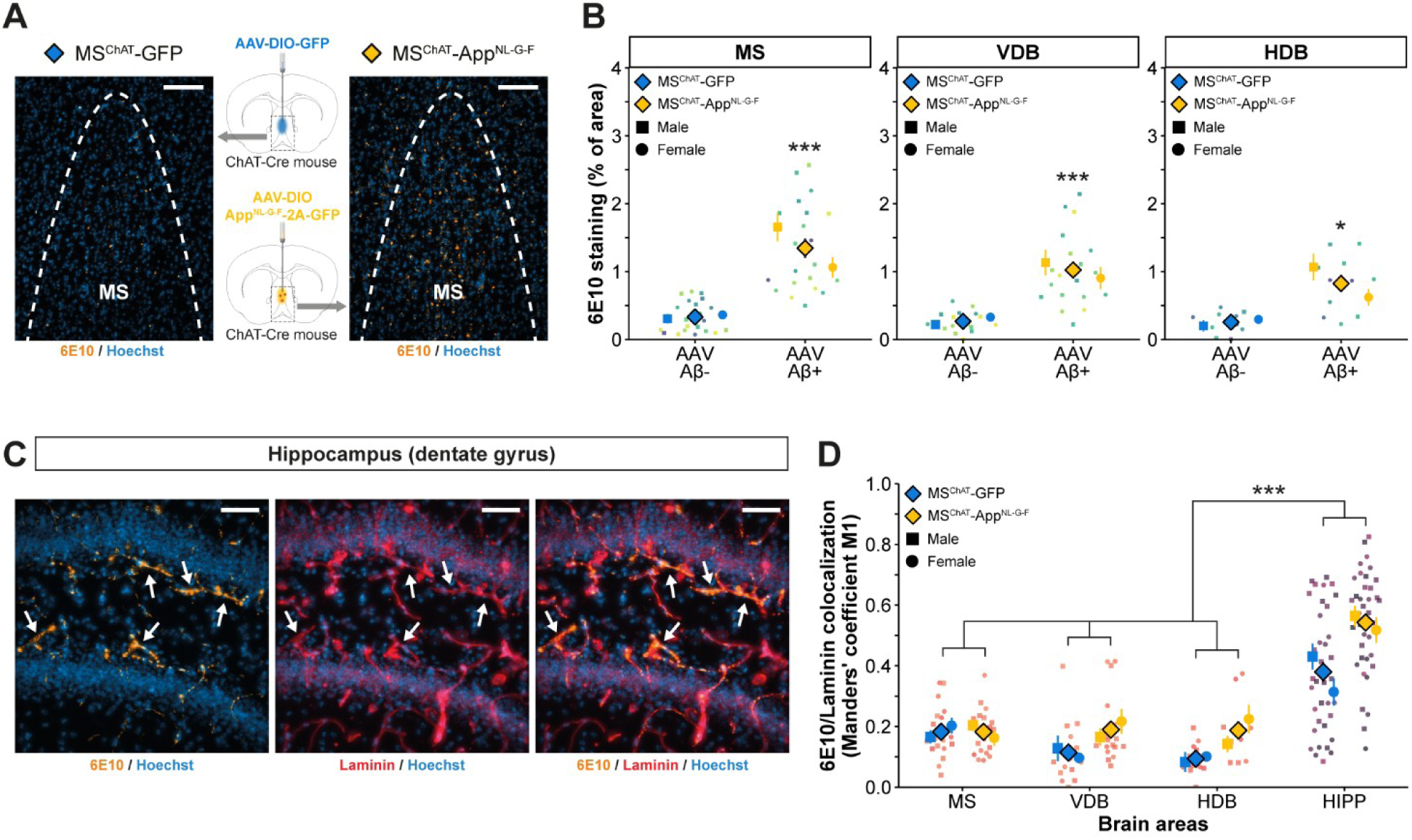
Enhanced colocalization between the 6E10 amyloid marker and cerebral vasculature outside the BF in *MS^ChAT^-App*^NL-G-F^ mice. (A) Coronal MS sections showing amyloid deposits (6E10, orange) and cell nuclei (Hoechst, blue) in *MS^ChAT^-GFP* (control; left) and *MS^ChAT^-App*^NL-G-F^ (right) mice (scale bar: 100 μm). (B) Quantification revealed significantly increased Aβ accumulation in the MS, VDB, and HDB of *MS^ChAT^-App*^NL-G-F^ mice compared with *MS^ChAT^-GFP* controls. See Figure 2B for gradient color-coding. (C) Representative hippocampal section from a *MS^ChAT^-App*^NL-G-F^ mouse show colocalization of amyloid deposits (6E10; orange) with blood vessels (laminin, red), highlighted by white arrows in the dentate gyrus (nuclei stained with Hoechst, blue) (scale bar: 50 μm). (D) Quantitative analysis using Mander’s coefficient (M1) showed that the proportion of 6E10 signal overlapping with laminin-positive vasculature was higher in the hippocampus than in the MS, VDB, and HDB in both *MS^ChAT^-GFP* and *MS^ChAT^-App*^NL-G-F^ mice. See Figure 1C for gradient color-coding. Statistical analysis: linear mixed-model ANOVA with Holm-Bonferroni correction for multiple pairwise comparisons where appropriate (*n* = 3 males/group, *n* = 3 females/group); mean ± SEM. Symbols: ▪ males, ● females, ♦ males and females, * *p* < 0.05, ** *p* < 0.01, *** *p* < 0.001.

**Figure S2.**
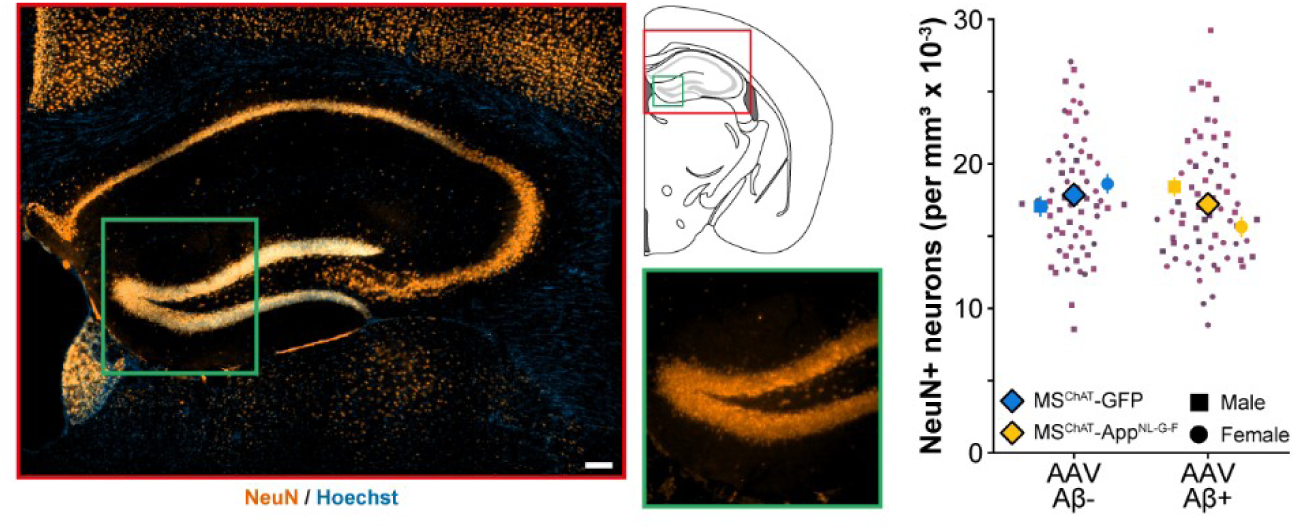
Broadcasted Aβ deposits into the hippocampus of *MS^ChAT^-App*^NL-G-F^ mice do not induce neuronal loss. Coronal hippocampal section showing the neuronal marker NeuN (orange) and cell nuclei (Hoechst, blue) (scale bar: 200 μm). Quantification of NeuN-positive cells revealed no difference between *MS^ChAT^-GFP* and *MS^ChAT^-App*^NL-G-F^ mice. Statistical analysis: linear mixed-model ANOVA with Holm-Bonferroni correction for multiple pairwise comparisons where appropriate (*n* = 4 males/group, *n* = 4 females/group); mean ± SEM. Symbols: ▪ males, ● females, ♦ males and females, * *p* < 0.05, ** *p* < 0.01, *** *p* < 0.001.

**Figure S3.**
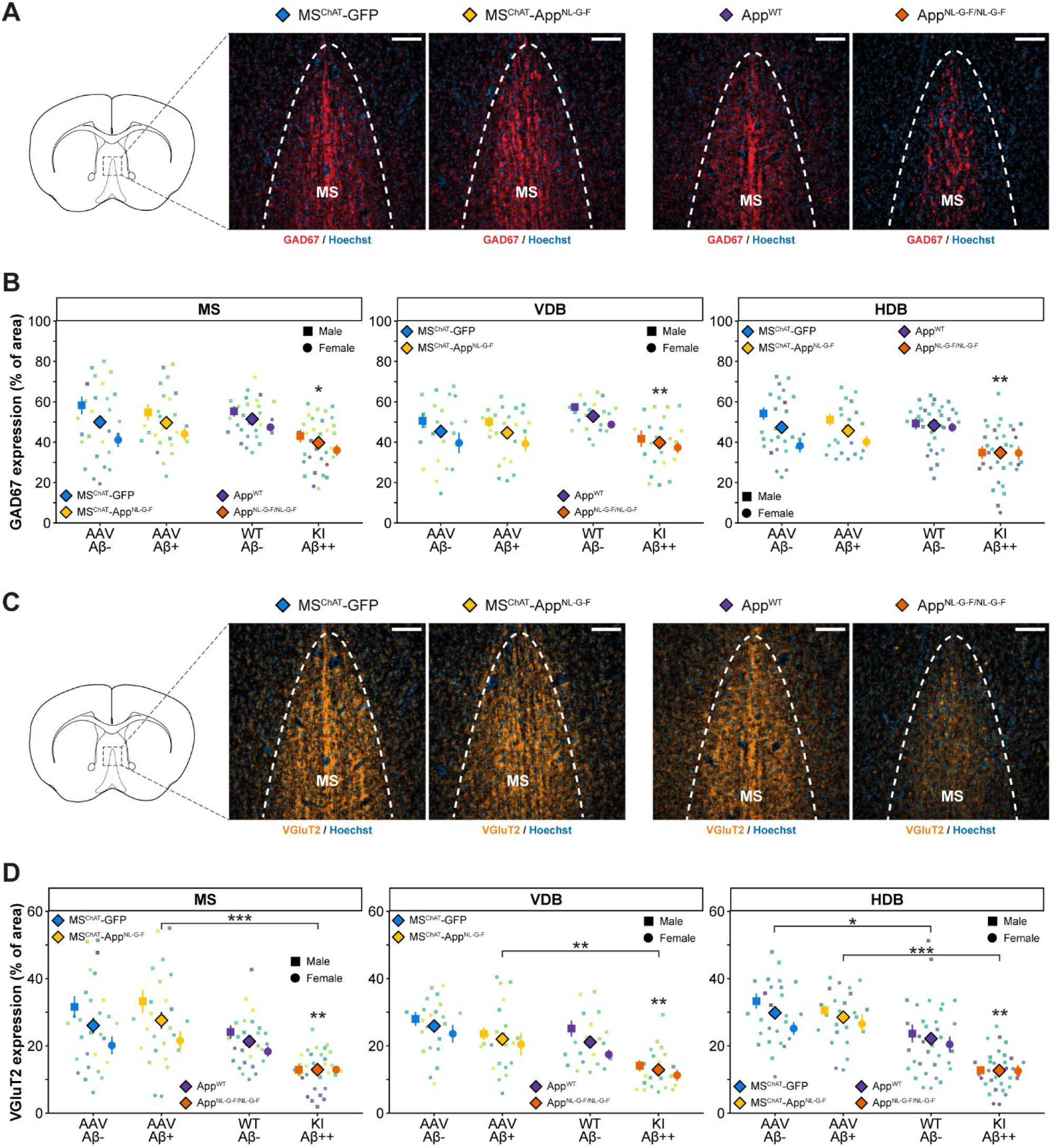
GABAergic and glutamatergic neurotransmission is preserved in *MS^ChAT^-App*^NL-G-F^ mice but impaired in *App*^NL-G-F/NL-G-F^ mice. (A) Coronal MS sections showing immunostaining for glutamic acid decarboxylase 67 (GAD67, red), and nuclei (Hoechst, blue) across genotypes from left to right: *MS^ChAT^-GFP* (control), *MS^ChAT^-App*^NL-G-F^, *App*^WT^ (control), and *App*^NL-G-F/NL-G-F^ mice (scale bar: 100 μm). (B) GAD67 expression in the MS, VDB, and HDB did not differ between *MS^ChAT^-GFP* and *MS^ChAT^-App*^NL-G-F^ mice. In contrast, *App*^NL-G-F/NL-G-F^ mice showed significantly reduced GAD67 expression across all three BF regions compared with *App^WT^* controls. See Figure 2B for gradient color-coding. (C) Coronal MS sections showing immunostaining for vesicular glutamate transporter 2 (VGluT2, orange) and nuclei (Hoechst, blue) across genotypes arranged from left to right: *MS^ChAT^-GFP* (control), *MS^ChAT^-App*^NL-G-F^, *App*^WT^ (control), and *App*^NL-G-F/NL-G-F^ mice (scale bar: 100 μm). (D) VGluT2 expression in the MS, VDB, and HDB did not differ between *MS^ChAT^-GFP* and *MS^ChAT^-App*^NL-G-F^ mice. In contrast, *App*^NL-G-F/NL-G-F^ mice exhibited significantly reduced VGluT2 expression in all BF regions compared with *App^WT^* controls. VGluT2 levels were also consistently lower in *App*^NL-G-F/NL-G-F^ mice than in *MS^ChAT^-App*^NL-G-F^ mice. See Figure 2B for gradient color-coding. Statistical analysis: linear mixed-model ANOVA with Holm-Bonferroni correction for multiple pairwise comparisons where appropriate (*n* = 4 males/group, *n* = 4 females/group); mean ± SEM. Asterisks on *MS^ChAT^-App*^NL-G-F^ and *App*^NL-G-F/NL-G-F^ groups denote significance relative to their corresponding controls. Symbols: ▪ males, ● females, ♦ males and females, * *p* < 0.05, ** *p* < 0.01, *** *p* < 0.001.

**Figure S4.**
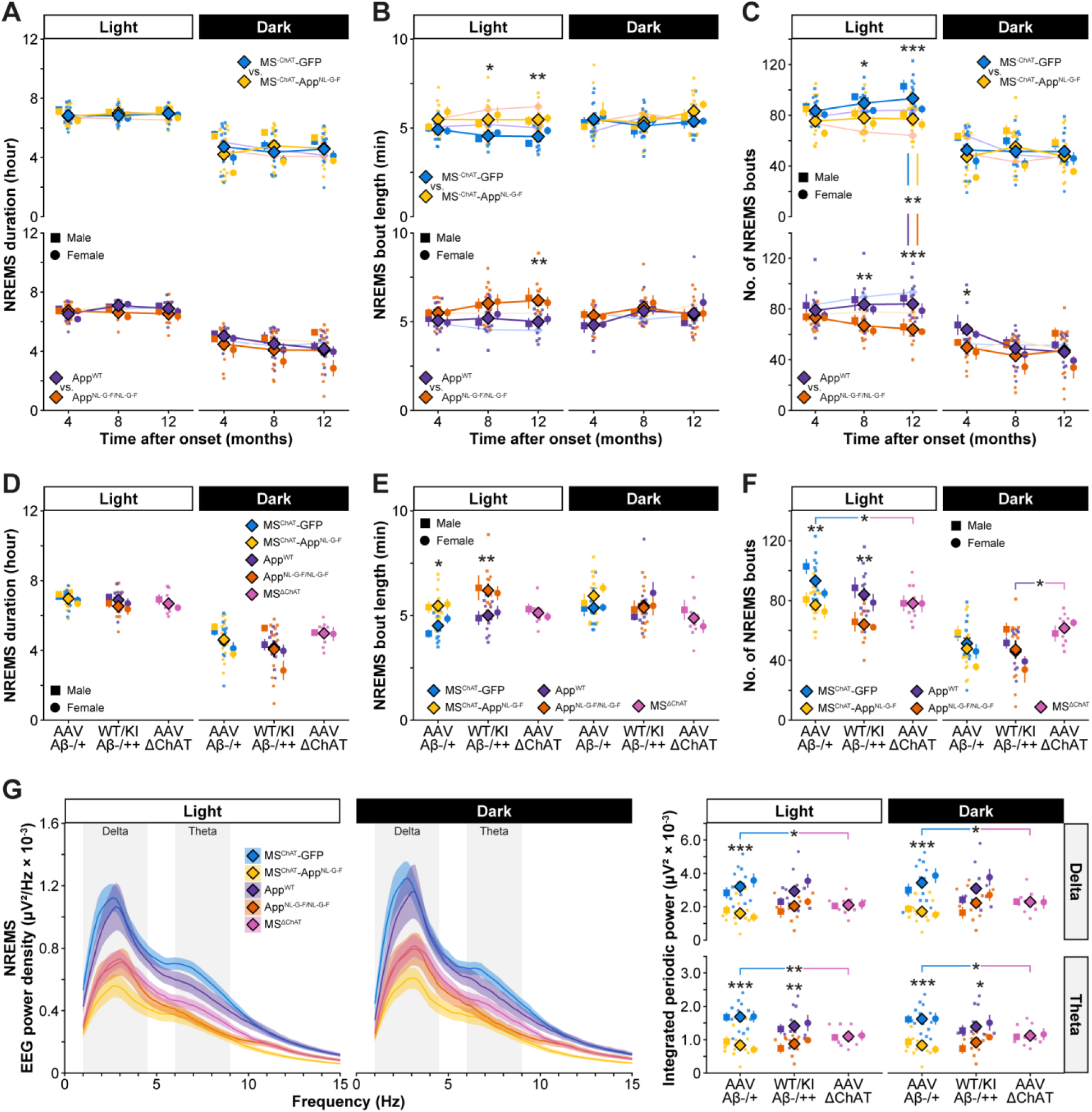
Shared alterations in NREMS phenotypes in *MS^ChAT^-App*^NL-G-F^, *App*^NL-G-F/NL-G-F^ and *MS^ΔChAT^* mice. (A) Across the one-year monitoring period, total NREMS duration remained comparable between groups during both light and dark phases. (B) During the light period, NREMS bout length was significantly increased in *MS^ChAT^-App*^NL-G-F^ mice at 8 and 12 months post-injection compared with *MS^ChAT^-GFP* controls. A similar increase was observed in *App*^NL-G-F/NL-G-F^ mice relative to *App^WT^* controls at 12 months, mirroring the phenotype seen in *MS^ChAT^-App*^NL-G-F^ animals. (C) *MS^ChAT^-GFP* mice showed an age-dependent increase in NREMS bout number during the light phase, whereas this effect was absent in *MS^ChAT^-App*^NL-G-F^ mice, resulting in significant differences at 8 and 12 months post-injection. In *App*^NL-G-F/NL-G-F^ mice, NREMS bout number was reduced compared with *App^WT^* controls at 12 months in the light period, with an additional reduction observed at 4 months during the dark phase. At 12 months of amyloid pathology, group differences were also evident in the light phase, with *MS^ChAT^-App*^NL-G-F^ and *MS^ChAT^-GFP* mice respectively differing from *App*^NL-G-F/NL-G-F^ and *App^WT^* controls. (D) Six months after AAV injection, *MS^ΔChAT^* mice did not differ in NREMS duration compared with other groups at 12 months of amyloid pathology. (E) NREMS bout length was comparable between *MS^ΔChAT^* mice and all other groups. (F) During the light phase, *MS^ΔChAT^* mice showed a reduced number of NREMS bouts compared with *MS^ChAT^-GFP* controls. In addition, during the dark phase, *MS^ΔChAT^* mice exhibited a higher number of NREMS bouts relative to *App^WT^* controls. (G) EEG spectral analysis (FOOOF) revealed decreased integrated periodic delta (1-4.4 Hz) and theta (6-9 Hz) power during both light and dark phases in *MS^ChAT^-App*^NL-G-F^ mice compared with *MS^ChAT^-GFP* controls. In *App*^NL-G-F/NL-G-F^ mice, only theta power was significantly reduced across both phases compared with *App^WT^* controls. Similar to the *MS^ChAT^-App*^NL-G-F^ model, *MS^ΔChAT^* mice also showed reduced delta and theta power in both light and dark phases relative to *MS^ChAT^-GFP* controls. Statistical analysis: linear mixed-model and standard ANOVAs with Holm-Bonferroni correction for multiple pairwise comparisons where appropriate (*n* = 5-8 males/group, *n* = 5-8 females/group); mean ± SEM. Symbols: ▪ males, ● females, ♦ males and females, * *p* < 0.05, ** *p* < 0.01, *** *p* < 0.001.

**Figure S5.**
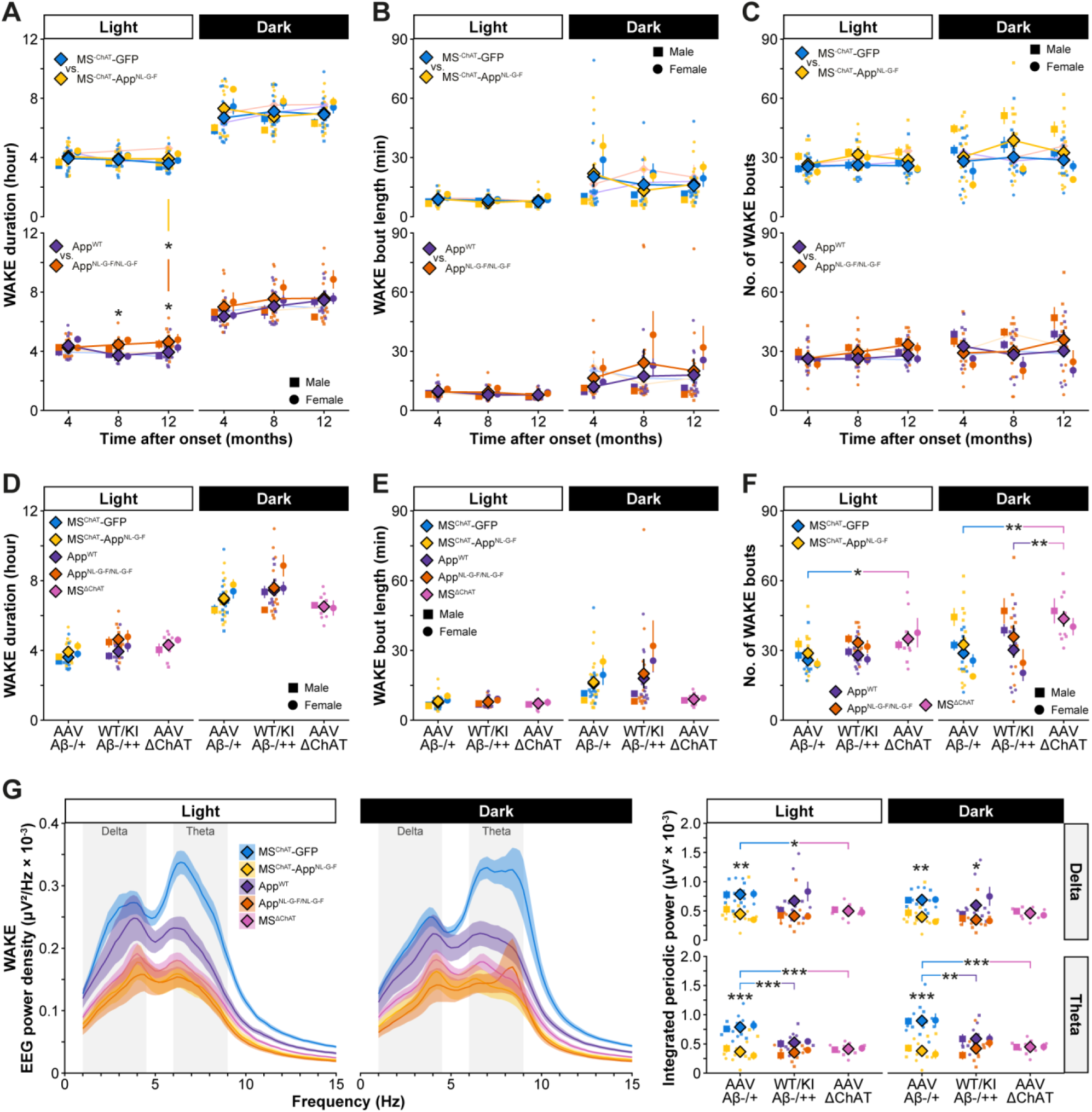
Differential wakefulness phenotypes in *MS^ChAT^-App*^NL-G-F^, *App*^NL-G-F/NL-G-F^ and *MS^ΔChAT^* mice. (A) Total wakefulness duration remained comparable between *MS^ChAT^-App*^NL-G-F^ and *MS^ChAT^-GFP* mice across the one-year recording period in both light and dark phases. In contrast, *App*^NL-G-F/NL-G-F^ mice exhibited increased wakefulness compared with *App^WT^* controls at 8 and 12 months during the light phase only. At 12 months of amyloid pathology, *App*^NL-G-F/NL-G-F^ mice also differed from *MS^ChAT^-App*^NL-G-F^ mice. (B) Wakefulness bout length remained stable and did not differ between *MS^ChAT^-App*^NL-G-F^ and *MS^ChAT^-GFP* mice, nor between *App*^NL-G-F/NL-G-F^ and *App^WT^* mice, across all time points and circadian phases. (C) The number of wakefulness bouts was also stable across time and did not differ between *MS^ChAT^-App*^NL-G-F^ and *MS^ChAT^-GFP* mice or between *App*^NL-G-F/NL-G-F^ and *App^WT^* controls, in both light and dark phases. (D) Six months after AAV injection, *MS^ΔChAT^* mice did not differ in total wakefulness duration compared with other groups at 12 months of amyloid pathology. (E) Wakefulness bout length was comparable between *MS^ΔChAT^* mice and all other groups. (F) *MS^ΔChAT^* mice exhibited an increased number of wakefulness bouts during both light and dark phases compared with *MS^ChAT^-GFP* controls. During the dark phase, this increase was also significant relative to *App^WT^* controls. (G) EEG spectral analysis (FOOOF) revealed decreased integrated periodic delta (1-4.4 Hz) and theta (6-9 Hz) power during both light and dark phases in *MS^ChAT^-App*^NL-G-F^ mice compared with *MS^ChAT^-GFP* controls. In *App*^NL-G-F/NL-G-F^ mice, delta power was reduced in the dark phase relative to *App^WT^* controls, while theta power was lower in *App^WT^* mice compared with *MS^ChAT^-GFP* in both phases. Similar to *MS^ChAT^-App*^NL-G-F^ mice, *MS^ΔChAT^* mice showed reduced theta power in both light and dark phases relative to *MS^ChAT^-GFP* controls, with a reduction in delta power restricted to the light phase. Statistical analysis: linear mixed-model and standard ANOVAs with Holm-Bonferroni correction for multiple pairwise comparisons where appropriate (*n* = 5-8 males/group, *n* = 5-8 females/group); mean ± SEM. Symbols: ▪ males, ● females, ♦ males and females, * *p* < 0.05, ** *p* < 0.01, *** *p* < 0.001.

**Figure S6.**
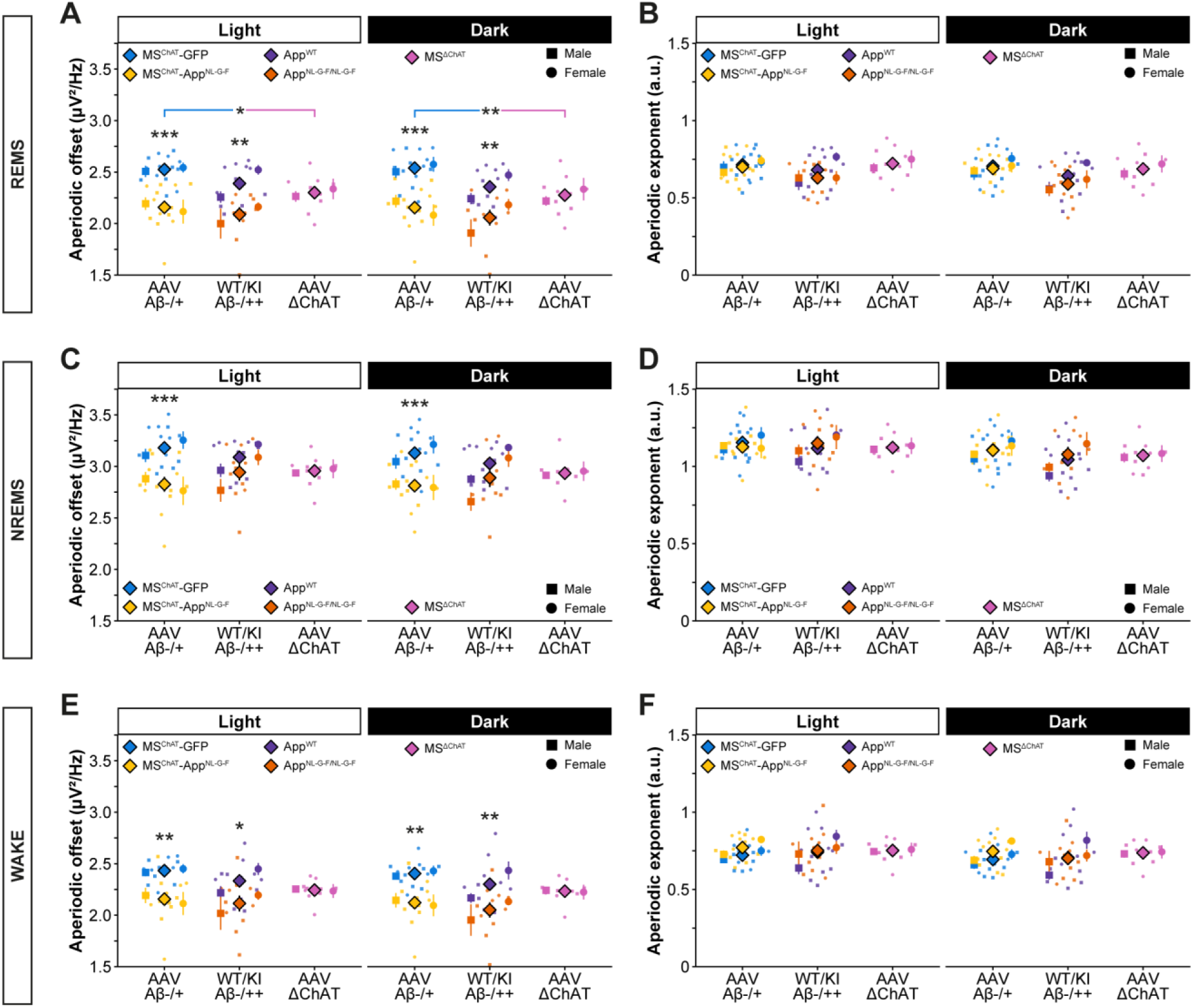
Aperiodic EEG features across vigilance states in *MS^ChAT^-App*^NL-G-F^, *App*^NL-G-F/NL-G-F^, and *MS^ΔChAT^* mice. (A) In REMS, the aperiodic offset was reduced in *MS^ChAT^-App*^NL-G-F^ and *App*^NL-G-F/NL-G-F^ mice compared with their respective controls during both light and dark phases. A similar reduction was observed in *MS^ΔChAT^* mice relative to *MS^ChAT^-GFP* controls. (B) The aperiodic exponent in REMS did not differ between groups. (C) In NREMS, the aperiodic offset was selectively reduced in *MS^ChAT^-App*^NL-G-F^ mice compared with *MS^ChAT^-GFP* controls in both light and dark phases. (D) The aperiodic exponent in NREMS remained unchanged across groups. (E) During wakefulness, the aperiodic offset was reduced in *MS^ChAT^-App*^NL-G-F^ and *App*^NL-G-F/NL-G-F^ mice relative to their respective controls in both phases, whereas *MS^ΔChAT^* mice did not differ from other groups. (F) The aperiodic exponent in wakefulness showed no group differences. Statistical analysis: standard ANOVA with Holm-Bonferroni correction for multiple pairwise comparisons where appropriate (*n* = 5-7 males/group, *n* = 5-7 females/group); mean ± SEM. Symbols: ▪ males, ● females, ♦ males and females, * *p* < 0.05, ** *p* < 0.01, *** *p* < 0.001.

**Figure S7.**
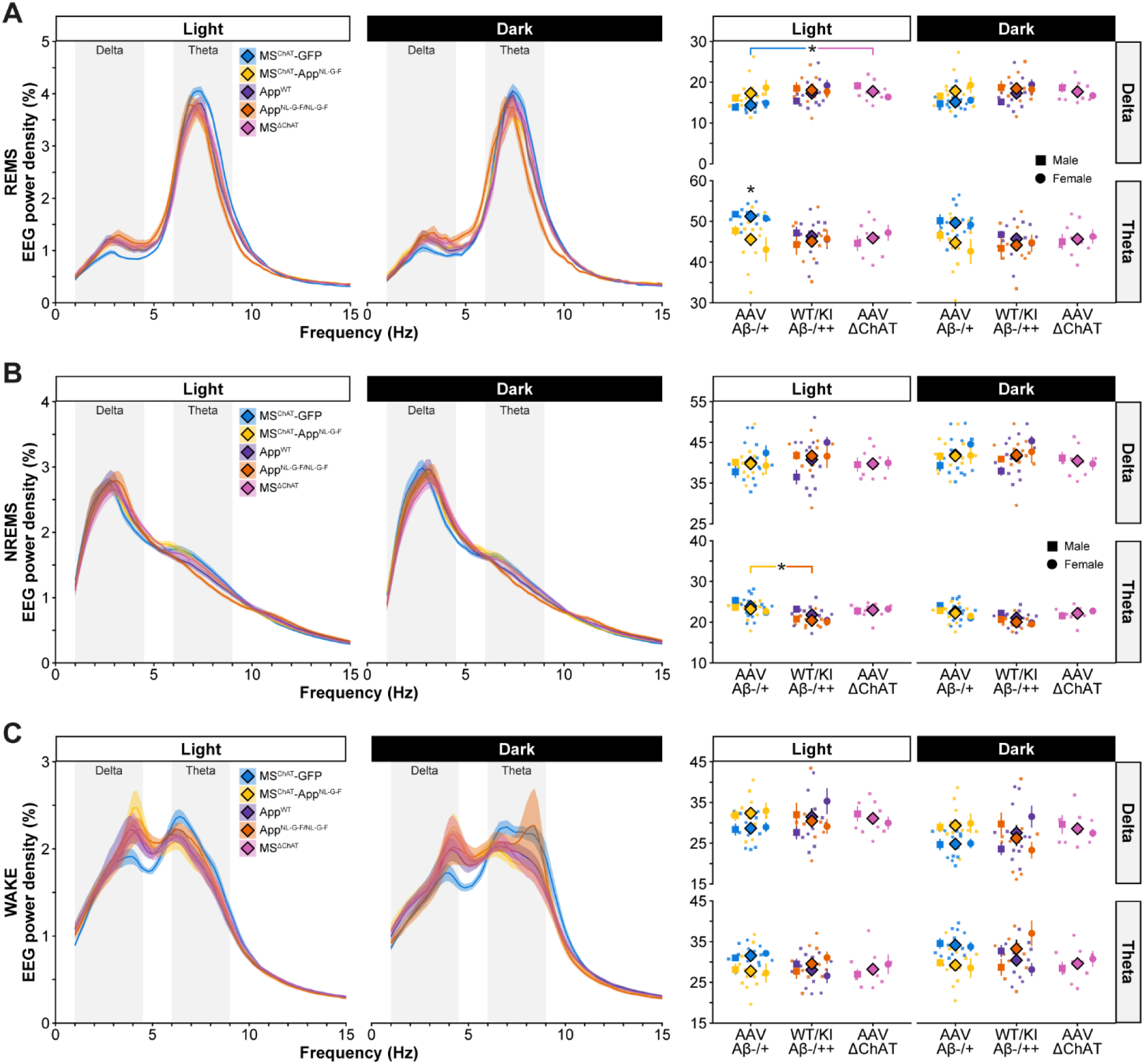
EEG spectral changes expressed as percentage of total power. (A) During REMS in the light phase, conventional EEG spectral analysis (expressed as percentage of total power) revealed a selective decrease in theta power in *MS^ChAT^-App*^NL-G-F^ mice compared with *MS^ChAT^-GFP* controls. In contrast, *MS^ΔChAT^* mice exhibited an increase in delta power relative to *MS^ChAT^-GFP* controls. (B) During NREMS in the light phase, theta power was significantly reduced in *App*^NL-G-F/NL-G-F^ mice compared with *MS^ChAT^-App*^NL-G-F^ animals. (C) During wakefulness, no significant spectral differences were detected between groups. Statistical analysis: ANOVA with Holm-Bonferroni correction for multiple pairwise comparisons where appropriate (*n* = 5-7 males/group, *n* = 5-7 females/group); mean ± SEM. Symbols: ▪ males, ● females, ♦ males and females, * *p* < 0.05, ** *p* < 0.01, *** *p* < 0.001.

## Notes

https://data.mendeley.com/preview/fdxrtbmfjk?a=d55f32c4-3622-4103-8ad9-88eabb17ecd9

### Competing Interest Statement

The authors have declared no competing interest.

### Summary of Updates

1. In the revised manuscript, we have reorganized the analyses and figures to now include direct statistical comparisons between the MSChAT-AppNL-G-F and AppNL-G-F/NL-G-F models throughout (as well as MS∆ChAT mice). This integrated approach confirms that both models display largely overlapping phenotypes, without altering the main conclusions of the study. 2. The newly introduced FOOOF analysis of the EEG spectrum (see revealed an even stronger reduction in theta power than previously observed using conventional normalized spectral approaches. 3. We now provide raw EEG traces (Figure 7B), which illustrate the differences between groups. These traces are complemented by integrated periodic power plots to provide a comprehensive view of the observed alterations. 4. We quantified the astrocytic marker GFAP in both the BF and hippocampus of MSChAT-AppNL-G-F and MSChAT-GFP mice, as well as in MSΔChAT mice in the BF. We found that the presence of amyloid in MSChAT-AppNL-G-F mice was associated with a significant increase in GFAP immunoreactivity in both regions, consistent with astrocytic activation in response to local amyloid pathology. In contrast, no such increase was observed in MSΔChAT mice, indicating that this glial response is specifically linked to amyloid deposition rather than cholinergic cell loss per se. These findings have now been incorporated into the manuscript, providing additional insight into the cellular specificity of amyloid-associated changes and highlighting a distinct glial component of the pathology. 5. MSChAT-AppNL-G-F mice also exhibited an increased number of epileptiform spikes relative to their respective controls, although to a lesser extent than KI mice. In contrast, MSΔChAT mice showed significantly fewer epileptiform spikes than MSChAT-App animals, suggesting that these modest epileptiform activities are primarily driven by amyloid pathology rather than by MS cholinergic neuron loss per se. These new analyses have now been included in the manuscript and are discussed in the context of the respective contributions of amyloid pathology and cholinergic loss, thereby providing a clearer understanding of the mechanisms underlying the observed phenotypes in our models 6. A new author has been added

## REFERENCES

1. Korczyn, A.D., and Grinberg, L.T. (2024). Is Alzheimer disease a disease? Nat Rev Neurol 20, 245–251. 10.1038/s41582-024-00940-4.

2. Wilson, D.M., 3rd, Cookson, M.R., Van Den Bosch, L., Zetterberg, H., Holtzman, D.M., and Dewachter, I. (2023). Hallmarks of neurodegenerative diseases. Cell 186, 693–714. 10.1016/j.cell.2022.12.032.

3. Hardy, J. (2025). Alzheimer’s Disease: Treatment Challenges for the Future. J Neurochem 169, e70176. 10.1111/jnc.70176.

4. Walsh, D.M., and Selkoe, D.J. (2020). Amyloid beta-protein and beyond: the path forward in Alzheimer’s disease. Curr Opin Neurobiol 61, 116–124. 10.1016/j.conb.2020.02.003.

5. Ehrenberg, A.J., Kelberman, M.A., Liu, K.Y., Dahl, M.J., Weinshenker, D., Falgàs, N., Dutt, S., Mather, M., Ludwig, M., Betts, M.J., et al. (2023). Priorities for research on neuromodulatory subcortical systems in Alzheimer’s disease: Position paper from the NSS PIA of ISTAART. Alzheimers & Dementia 19, 2182–2196. 10.1002/alz.12937.

6. Ananth, M.R., Rajebhosale, P., Kim, R., Talmage, D.A., and Role, L.W. (2023). Basal forebrain cholinergic signalling: development, connectivity and roles in cognition. Nat Rev Neurosci 24, 233–251. 10.1038/s41583-023-00677-x.

7. Picciotto, M.R., Higley, M.J., and Mineur, Y.S. (2012). Acetylcholine as a neuromodulator: cholinergic signaling shapes nervous system function and behavior. Neuron 76, 116–129. 10.1016/j.neuron.2012.08.036.

8. Xu, M., Chung, S., Zhang, S., Zhong, P., Ma, C., Chang, W.C., Weissbourd, B., Sakai, N., Luo, L., Nishino, S., and Dan, Y. (2015). Basal forebrain circuit for sleep-wake control. Nat Neurosci 18, 1641–1647. 10.1038/nn.4143.

9. Irmak, S.O., and de Lecea, L. (2014). Basal forebrain cholinergic modulation of sleep transitions. Sleep 37, 1941–1951. 10.5665/sleep.4246.

10. Galvin, V.C., Arnsten, A.F.T., and Wang, M. (2018). Evolution in Neuromodulation-The Differential Roles of Acetylcholine in Higher Order Association vs. Primary Visual Cortices. Front Neural Circuits 12, 67. 10.3389/fncir.2018.00067.

11. Desikan, S., Koser, D.E., Neitz, A., and Monyer, H. (2018). Target selectivity of septal cholinergic neurons in the medial and lateral entorhinal cortex. Proc Natl Acad Sci U S A 115, E2644–E2652. 10.1073/pnas.1716531115.

12. Dobryakova, Y.V., Bolshakov, A.P., Korotkova, T., and Rozov, A.V. (2025). Acetylcholine in the hippocampus: problems and achievements. Front Neural Circuits 19, 1491820. 10.3389/fncir.2025.1491820.

13. Orlando, I.F., Shine, J.M., Robbins, T.W., Rowe, J.B., and O’Callaghan, C. (2023). Noradrenergic and cholinergic systems take centre stage in neuropsychiatric diseases of ageing. Neurosci Biobehav Rev 149, 105167. 10.1016/j.neubiorev.2023.105167.

14. Li, B., Ma, C., Huang, Y.A., Ding, X., Silverman, D., Chen, C., Darmohray, D., Lu, L., Liu, S., Montaldo, G., et al. (2023). Circuit mechanism for suppression of frontal cortical ignition during NREM sleep. Cell 186, 5739–5750 e5717. 10.1016/j.cell.2023.11.012.

15. Zaborszky, L., Gombkoto, P., Varsanyi, P., Gielow, M.R., Poe, G., Role, L.W., Ananth, M., Rajebhosale, P., Talmage, D.A., Hasselmo, M.E., et al. (2018). Specific Basal Forebrain-Cortical Cholinergic Circuits Coordinate Cognitive Operations. J Neurosci 38, 9446–9458. 10.1523/JNEUROSCI.1676-18.2018.

16. Li, X., Yu, B., Sun, Q., Zhang, Y., Ren, M., Zhang, X., Li, A., Yuan, J., Madisen, L., Luo, Q., et al. (2018). Generation of a whole-brain atlas for the cholinergic system and mesoscopic projectome analysis of basal forebrain cholinergic neurons. Proc Natl Acad Sci U S A 115, 415–420. 10.1073/pnas.1703601115.

17. Davies, P., and Maloney, A.J. (1976). Selective loss of central cholinergic neurons in Alzheimer’s disease. Lancet 2, 1403. 10.1016/s0140-6736(76)91936-x.

18. Bartus, R.T., Dean, R.L., 3rd, Beer, B., and Lippa, A.S. (1982). The cholinergic hypothesis of geriatric memory dysfunction. Science 217, 408–414. 10.1126/science.7046051.

19. Rossor, M.N., Garrett, N.J., Johnson, A.L., Mountjoy, C.Q., Roth, M., and Iversen, L.L. (1982). A post-mortem study of the cholinergic and GABA systems in senile dementia. Brain 105, 313–330. 10.1093/brain/105.2.313.

20. Whitehouse, P.J., Price, D.L., Struble, R.G., Clark, A.W., Coyle, J.T., and Delon, M.R. (1982). Alzheimer’s disease and senile dementia: loss of neurons in the basal forebrain. Science 215, 1237–1239. 10.1126/science.7058341.

21. Geula, C., Dunlop, S.R., Ayala, I., Kawles, A.S., Flanagan, M.E., Gefen, T., and Mesulam, M.M. (2021). Basal forebrain cholinergic system in the dementias: Vulnerability, resilience, and resistance. J Neurochem 158, 1394–1411. 10.1111/jnc.15471.

22. Aghourian, M., Aumont, E., Grothe, M.J., Soucy, J.P., Rosa-Neto, P., and Bedard, M.A. (2021). FEOBV-PET to quantify cortical cholinergic denervation in AD: Relationship to basal forebrain volumetry. J Neuroimaging 31, 1077–1081. 10.1111/jon.12921.

23. Kilimann, I., Grothe, M., Heinsen, H., Alho, E.J., Grinberg, L., Amaro, E., Jr., Dos Santos, G.A., da Silva, R.E., Mitchell, A.J., Frisoni, G.B., et al. (2014). Subregional basal forebrain atrophy in Alzheimer’s disease: a multicenter study. J Alzheimers Dis 40, 687–700. 10.3233/JAD-132345.

24. Berry, A.S., and Harrison, T.M. (2023). New perspectives on the basal forebrain cholinergic system in Alzheimer’s disease. Neurosci Biobehav Rev 150, 105192. 10.1016/j.neubiorev.2023.105192.

25. Hampel, H., Mesulam, M.M., Cuello, A.C., Khachaturian, A.S., Vergallo, A., Farlow, M.R., Snyder, P.J., Giacobini, E., and Khachaturian, Z.S. (2019). Revisiting the Cholinergic Hypothesis in Alzheimer’s Disease: Emerging Evidence from Translational and Clinical Research. J Prev Alzheimers Dis 6, 2–15. 10.14283/jpad.2018.43.

26. Grothe, M., Zaborszky, L., Atienza, M., Gil-Neciga, E., Rodriguez-Romero, R., Teipel, S.J., Amunts, K., Suarez-Gonzalez, A., and Cantero, J.L. (2010). Reduction of basal forebrain cholinergic system parallels cognitive impairment in patients at high risk of developing Alzheimer’s disease. Cereb Cortex 20, 1685–1695. 10.1093/cercor/bhp232.

27. Zheng, W., Li, H., Cui, B., Liang, P., Wu, Y., Han, X., Li, C.R., Li, K., and Wang, Z. (2020). Altered multimodal magnetic resonance parameters of basal nucleus of Meynert in Alzheimer’s disease. Ann Clin Transl Neurol 7, 1919–1929. 10.1002/acn3.51176.

28. Schmitz, T.W., Mur, M., Aghourian, M., Bedard, M.A., Spreng, R.N., and Alzheimer’s Disease Neuroimaging, I. (2018). Longitudinal Alzheimer’s Degeneration Reflects the Spatial Topography of Cholinergic Basal Forebrain Projections. Cell Rep 24, 38–46. 10.1016/j.celrep.2018.06.001.

29. Ballinger, E.C., Ananth, M., Talmage, D.A., and Role, L.W. (2016). Basal Forebrain Cholinergic Circuits and Signaling in Cognition and Cognitive Decline. Neuron 91, 1199–1218. 10.1016/j.neuron.2016.09.006.

30. Taza, M., Schmitz, T.W., and Spreng, R.N. (2025). Structural changes to the basal forebrain cholinergic system in the continuum of Alzheimer disease. Handb Clin Neurol 211, 81–93. 10.1016/B978-0-443-19088-9.00013-5.

31. Schmitz, T.W., Nathan Spreng, R., and Alzheimer’s Disease Neuroimaging, I. (2016). Basal forebrain degeneration precedes and predicts the cortical spread of Alzheimer’s pathology. Nat Commun 7, 13249. 10.1038/ncomms13249.

32. Theofilas, P., Dunlop, S., Heinsen, H., and Grinberg, L.T. (2015). Turning on the Light Within: Subcortical Nuclei of the Isodentritic Core and their Role in Alzheimer’s Disease Pathogenesis. Journal of Alzheimers Disease 46, 17–34. 10.3233/Jad-142682.

33. Bartus, R.T. (2000). On neurodegenerative diseases, models, and treatment strategies: lessons learned and lessons forgotten a generation following the cholinergic hypothesis. Exp Neurol 163, 495–529. 10.1006/exnr.2000.7397.

34. Nollet, M., Franks, N.P., and Wisden, W. (2023). Understanding Sleep Regulation in Normal and Pathological Conditions, and Why It Matters. J Huntingtons Dis 12, 105–119. 10.3233/JHD-230564.

35. Sulaman, B.A., Wang, S., Tyan, J., and Eban-Rothschild, A. (2023). Neuro-orchestration of sleep and wakefulness. Nat Neurosci 26, 196–212. 10.1038/s41593-022-01236-w.

36. Niwa, Y., Kanda, G.N., Yamada, R.G., Shi, S., Sunagawa, G.A., Ukai-Tadenuma, M., Fujishima, H., Matsumoto, N., Masumoto, K.H., Nagano, M., et al. (2018). Muscarinic Acetylcholine Receptors Chrm1 and Chrm3 Are Essential for REM Sleep. Cell Rep 24, 2231–2247 e2237. 10.1016/j.celrep.2018.07.082.

37. Pase, M.P., Himali, J.J., Grima, N.A., Beiser, A.S., Satizabal, C.L., Aparicio, H.J., Thomas, R.J., Gottlieb, D.J., Auerbach, S.H., and Seshadri, S. (2017). Sleep architecture and the risk of incident dementia in the community. Neurology 89, 1244–1250. 10.1212/WNL.0000000000004373.

38. Taillard, J., Sagaspe, P., Berthomier, C., Brandewinder, M., Amieva, H., Dartigues, J.F., Rainfray, M., Harston, S., Micoulaud-Franchi, J.A., and Philip, P. (2019). Non-REM Sleep Characteristics Predict Early Cognitive Impairment in an Aging Population. Front Neurol 10, 197. 10.3389/fneur.2019.00197.

39. André, C., Martineau-Dussault, M.E., Daneault, V., Blais, H., Frenette, S., Lorrain, D., Hudon, C., Bastien, C., Petit, D., Lafrenière, A., et al. (2023). REM sleep is associated with the volume of the cholinergic basal forebrain in aMCI individuals. Alzheimers Research & Therapy 15. ARTN 151 10.1186/s13195-023-01265-y.

40. Leary, E.B., Watson, K.T., Ancoli-Israel, S., Redline, S., Yaffe, K., Ravelo, L.A., Peppard, P.E., Zou, J., Goodman, S.N., Mignot, E., and Stone, K.L. (2020). Association of Rapid Eye Movement Sleep With Mortality in Middle-aged and Older Adults. JAMA Neurol 77, 1241–1251. 10.1001/jamaneurol.2020.2108.

41. Della Monica, C., Johnsen, S., Atzori, G., Groeger, J.A., and Dijk, D.J. (2018). Rapid Eye Movement Sleep, Sleep Continuity and Slow Wave Sleep as Predictors of Cognition, Mood, and Subjective Sleep Quality in Healthy Men and Women, Aged 20-84 Years. Front Psychiatry 9, 255. 10.3389/fpsyt.2018.00255.

42. Djonlagic, I., Mariani, S., Fitzpatrick, A.L., Van Der Klei, V., Johnson, D.A., Wood, A.C., Seeman, T., Nguyen, H.T., Prerau, M.J., Luchsinger, J.A., et al. (2021). Macro and micro sleep architecture and cognitive performance in older adults. Nat Hum Behav 5, 123–145. 10.1038/s41562-020-00964-y.

43. Scullin, M.K., Gao, C., Fillmore, P., Roberts, R.L., Pruett, N., and Bliwise, D.L. (2019). Rapid eye movement sleep mediates age-related decline in prospective memory consolidation. Sleep 42. 10.1093/sleep/zsz055.

44. Ba, W., Nollet, M., Yin, C., Yu, X., Wong, S., Miao, A., Beckwith, E.J., Harding, E.C., Ma, Y., Yustos, R., et al. (2024). A REM-active basal ganglia circuit that regulates anxiety. Curr Biol 34, 3301–3314 e3304. 10.1016/j.cub.2024.06.010.

45. Nollet, M., Hicks, H., McCarthy, A.P., Wu, H., Moller-Levet, C.S., Laing, E.E., Malki, K., Lawless, N., Wafford, K.A., Dijk, D.J., and Winsky-Sommerer, R. (2019). REM sleep’s unique associations with corticosterone regulation, apoptotic pathways, and behavior in chronic stress in mice. Proc Natl Acad Sci U S A 116, 2733–2742. 10.1073/pnas.1816456116.

46. Yu, X., Nollet, M., Franks, N.P., and Wisden, W. (2025). Sleep and the recovery from stress. Neuron. 10.1016/j.neuron.2025.04.028.

47. Chetty, C.A., Bhardwaj, H., Kumar, G.P., Devanand, T., Sekhar, C.S.A., Akturk, T., Kiyi, I., Yener, G., Guntekin, B., Joseph, J., and Adaikkan, C. (2024). EEG biomarkers in Alzheimer’s and prodromal Alzheimer’s: a comprehensive analysis of spectral and connectivity features. Alzheimers Res Ther 16, 236. 10.1186/s13195-024-01582-w.

48. Zawislak-Fornagiel, K., Ledwon, D., Bugdol, M., Grazynska, A., Slot, M., Tabaka-Pradela, J., Bieniek, I., and Siuda, J. (2024). Quantitative EEG Spectral and Connectivity Analysis for Cognitive Decline in Amnestic Mild Cognitive Impairment. J Alzheimers Dis 97, 1235–1247. 10.3233/JAD-230485.

49. Maezono, S.E.B., Kanuka, M., Tatsuzawa, C., Morita, M., Kawano, T., Kashiwagi, M., Nondhalee, P., Sakaguchi, M., Saito, T., Saido, T.C., and Hayashi, Y. (2020). Progressive Changes in Sleep and Its Relations to Amyloid-beta Distribution and Learning in Single App Knock-In Mice. eNeuro 7. 10.1523/ENEURO.0093-20.2020.

50. Pulver, R.L., Kronberg, E., Medenblik, L.M., Kheyfets, V.O., Ramos, A.R., Holtzman, D.M., Morris, J.C., Toedebusch, C.D., Sillau, S.H., Bettcher, B.M., et al. (2024). Mapping sleep’s oscillatory events as a biomarker of Alzheimer’s disease. Alzheimers Dement 20, 301–315. 10.1002/alz.13420.

51. Olgun, Y., Aksoy Poyraz, C., Bozluolcay, M., and Poyraz, B.C. (2024). Quantitative EEG in the Differential Diagnosis of Dementia Subtypes. J Geriatr Psychiatry Neurol 37, 368–378. 10.1177/08919887241227410.

52. Meghdadi, A.H., Salat, D., Hamilton, J., Hong, Y., Boeve, B.F., St Louis, E.K., Verma, A., and Berka, C. (2024). EEG and ERP biosignatures of mild cognitive impairment for longitudinal monitoring of early cognitive decline in Alzheimer’s disease. PLoS One 19, e0308137. 10.1371/journal.pone.0308137.

53. Kent, B.A., Strittmatter, S.M., and Nygaard, H.B. (2018). Sleep and EEG Power Spectral Analysis in Three Transgenic Mouse Models of Alzheimer’s Disease: APP/PS1, 3xTgAD, and Tg2576. J Alzheimers Dis 64, 1325–1336. 10.3233/JAD-180260.

54. Azami, H., Zrenner, C., Brooks, H., Zomorrodi, R., Blumberger, D.M., Fischer, C.E., Flint, A., Herrmann, N., Kumar, S., Lanctot, K., et al. (2023). Beta to theta power ratio in EEG periodic components as a potential biomarker in mild cognitive impairment and Alzheimer’s dementia. Alzheimers Res Ther 15, 133. 10.1186/s13195-023-01280-z.

55. Yuan, W., Zhi, W., Ma, L., Hu, X., Wang, Q., Zou, Y., and Wang, L. (2023). Neural Oscillation Disorder in the Hippocampal CA1 Region of Different Alzheimer’s Disease Mice. Curr Alzheimer Res 20, 350–359. 10.2174/1567205020666230808122643.

56. Chen, X., Li, Y., Li, R., Yuan, X., Liu, M., Zhang, W., and Li, Y. (2023). Multiple cross-frequency coupling analysis of resting-state EEG in patients with mild cognitive impairment and Alzheimer’s disease. Front Aging Neurosci 15, 1142085. 10.3389/fnagi.2023.1142085.

57. Scheijbeler, E.P., de Haan, W., Stam, C.J., Twisk, J.W.R., and Gouw, A.A. (2023). Longitudinal resting-state EEG in amyloid-positive patients along the Alzheimer’s disease continuum: considerations for clinical trials. Alzheimers Res Ther 15, 182. 10.1186/s13195-023-01327-1.

58. Kopcanova, M., Tait, L., Donoghue, T., Stothart, G., Smith, L., Flores-Sandoval, A.A., Davila-Perez, P., Buss, S., Shafi, M.M., Pascual-Leone, A., et al. (2024). Resting-state EEG signatures of Alzheimer’s disease are driven by periodic but not aperiodic changes. Neurobiol Dis 190, 106380. 10.1016/j.nbd.2023.106380.

59. Wang, S., Li, K., Zhao, S., Zhang, X., Yang, Z., Zhang, J., and Zhang, T. (2020). Early-stage dysfunction of hippocampal theta and gamma oscillations and its modulation of neural network in a transgenic 5xFAD mouse model. Neurobiol Aging 94, 121–129. 10.1016/j.neurobiolaging.2020.05.002.

60. Altunkaya, A., Deichsel, C., Kreuzer, M., Nguyen, D.M., Wintergerst, A.M., Rammes, G., Schneider, G., and Fenzl, T. (2024). Altered sleep behavior strengthens face validity in the ArcAbeta mouse model for Alzheimer’s disease. Sci Rep 14, 951. 10.1038/s41598-024-51560-3.

61. Lam, A.K.F., Carrick, J., Kao, C.H., Phillips, C.L., Zheng, Y.Z., Yee, B.J., Kim, J.W., Grunstein, R.R., Naismith, S.L., and D’Rozario, A.L. (2024). EEG slowing during REM sleep in older adults with subjective cognitive impairment and mild cognitive impairment. Sleep. 10.1093/sleep/zsae051.

62. Baker-Nigh, A., Vahedi, S., Davis, E.G., Weintraub, S., Bigio, E.H., Klein, W.L., and Geula, C. (2015). Neuronal amyloid-beta accumulation within cholinergic basal forebrain in ageing and Alzheimer’s disease. Brain 138, 1722–1737. 10.1093/brain/awv024.

63. Alldred, M.J., Penikalapati, S.C., Lee, S.H., Heguy, A., Roussos, P., and Ginsberg, S.D. (2021). Profiling Basal Forebrain Cholinergic Neurons Reveals a Molecular Basis for Vulnerability Within the Ts65Dn Model of Down Syndrome and Alzheimer’s Disease. Mol Neurobiol 58, 5141–5162. 10.1007/s12035-021-02453-3.

64. Kerbler, G.M., Fripp, J., Rowe, C.C., Villemagne, V.L., Salvado, O., Rose, S., Coulson, E.J., and Alzheimer’s Disease Neuroimaging, I. (2015). Basal forebrain atrophy correlates with amyloid beta burden in Alzheimer’s disease. Neuroimage Clin 7, 105–113. 10.1016/j.nicl.2014.11.015.

65. An, S., Sun, H., Wu, M., Xie, D., Hu, S.W., Ding, H.L., and Cao, J.L. (2021). Medial septum glutamatergic neurons control wakefulness through a septo-hypothalamic circuit. Curr Biol 31, 1379–1392 e1374. 10.1016/j.cub.2021.01.019.

66. Joshi, A., Salib, M., Viney, T.J., Dupret, D., and Somogyi, P. (2017). Behavior-Dependent Activity and Synaptic Organization of Septo-hippocampal GABAergic Neurons Selectively Targeting the Hippocampal CA3 Area. Neuron 96, 1342–1357 e1345. 10.1016/j.neuron.2017.10.033.

67. Salib, M., Joshi, A., Katona, L., Howarth, M., Micklem, B.R., Somogyi, P., and Viney, T.J. (2019). GABAergic Medial Septal Neurons with Low-Rhythmic Firing Innervating the Dentate Gyrus and Hippocampal Area CA3. J Neurosci 39, 4527–4549. 10.1523/JNEUROSCI.3024-18.2019.

68. Boyce, R., Glasgow, S.D., Williams, S., and Adamantidis, A. (2016). Causal evidence for the role of REM sleep theta rhythm in contextual memory consolidation. Science 352, 812–816. 10.1126/science.aad5252.

69. Colom, L.V., Castaneda, M.T., Banuelos, C., Puras, G., Garcia-Hernandez, A., Hernandez, S., Mounsey, S., Benavidez, J., and Lehker, C. (2010). Medial septal beta-amyloid 1-40 injections alter septo-hippocampal anatomy and function. Neurobiol Aging 31, 46–57. 10.1016/j.neurobiolaging.2008.05.006.

70. Wu, M.N., Kong, L.L., Zhang, J., Hu, M.M., Wang, Z.J., Cai, H.Y., and Qi, J.S. (2018). Amyloid beta protein injection into medial septum impairs hippocampal long-term potentiation and cognitive behaviors in rats. Sheng Li Xue Bao 70, 217–227.

71. Saito, T., Matsuba, Y., Mihira, N., Takano, J., Nilsson, P., Itohara, S., Iwata, N., and Saido, T.C. (2014). Single App knock-in mouse models of Alzheimer’s disease. Nat Neurosci 17, 661–663. 10.1038/nn.3697.

72. Donoghue, T., Haller, M., Peterson, E.J., Varma, P., Sebastian, P., Gao, R., Noto, T., Lara, A.H., Wallis, J.D., Knight, R.T., et al. (2020). Parameterizing neural power spectra into periodic and aperiodic components. Nature Neuroscience 23, 1655–U1288. 10.1038/s41593-020-00744-x.

73. Ameen, M.S., Jacobs, J., Schabus, M., Hoedlmoser, K., and Donoghue, T. (2025). Temporally resolved analyses of aperiodic features track neural dynamics during sleep. Commun Psychol 3, 160. 10.1038/s44271-025-00334-2.

74. Palop, J.J., Chin, J., Roberson, E.D., Wang, J., Thwin, M.T., Bien-Ly, N., Yoo, J., Ho, K.O., Yu, G.Q., Kreitzer, A., et al. (2007). Aberrant excitatory neuronal activity and compensatory remodeling of inhibitory hippocampal circuits in mouse models of Alzheimer’s disease. Neuron 55, 697–711. 10.1016/j.neuron.2007.07.025.

75. Minkeviciene, R., Rheims, S., Dobszay, M.B., Zilberter, M., Hartikainen, J., Fulop, L., Penke, B., Zilberter, Y., Harkany, T., Pitkanen, A., and Tanila, H. (2009). Amyloid beta-induced neuronal hyperexcitability triggers progressive epilepsy. J Neurosci 29, 3453–3462. 10.1523/JNEUROSCI.5215-08.2009.

76. Busche, M.A., Eichhoff, G., Adelsberger, H., Abramowski, D., Wiederhold, K.H., Haass, C., Staufenbiel, M., Konnerth, A., and Garaschuk, O. (2008). Clusters of hyperactive neurons near amyloid plaques in a mouse model of Alzheimer’s disease. Science 321, 1686–1689. 10.1126/science.1162844.

77. Busche, M.A., Chen, X., Henning, H.A., Reichwald, J., Staufenbiel, M., Sakmann, B., and Konnerth, A. (2012). Critical role of soluble amyloid-beta for early hippocampal hyperactivity in a mouse model of Alzheimer’s disease. Proc Natl Acad Sci U S A 109, 8740–8745. 10.1073/pnas.1206171109.

78. Mandybur, T.I., and Chuirazzi, C.C. (1990). Astrocytes and the plaques of Alzheimer’s disease. Neurology 40, 635–639. 10.1212/wnl.40.4.635.

79. Kamphuis, W., Middeldorp, J., Kooijman, L., Sluijs, J.A., Kooi, E.J., Moeton, M., Freriks, M., Mizee, M.R., and Hol, E.M. (2014). Glial fibrillary acidic protein isoform expression in plaque related astrogliosis in Alzheimer’s disease. Neurobiol Aging 35, 492–510. 10.1016/j.neurobiolaging.2013.09.035.

80. Pang, K., Jiang, R., Zhang, W., Yang, Z., Li, L.L., Shimozawa, M., Tambaro, S., Mayer, J., Zhang, B., Li, M., et al. (2022). An App knock-in rat model for Alzheimer’s disease exhibiting Abeta and tau pathologies, neuronal death and cognitive impairments. Cell Res 32, 157–175. 10.1038/s41422-021-00582-x.

81. Liu, H., Mei, F., Ye, R., Han, X., Wang, S., Ding, Y., Zhi, Y., Pang, K., Guo, W., and Lu, B. (2024). APOE3ch alleviates Abeta and tau pathology and neurodegeneration in the human APP(NL-G-F) cerebral organoid model of Alzheimer’s disease. Cell Res 34, 451–454. 10.1038/s41422-024-00957-w.

82. Gail Canter, R., Huang, W.C., Choi, H., Wang, J., Ashley Watson, L., Yao, C.G., Abdurrob, F., Bousleiman, S.M., Young, J.Z., Bennett, D.A., et al. (2019). 3D mapping reveals network-specific amyloid progression and subcortical susceptibility in mice. Commun Biol 2, 360. 10.1038/s42003-019-0599-8.

83. Cirrito, J.R., Yamada, K.A., Finn, M.B., Sloviter, R.S., Bales, K.R., May, P.C., Schoepp, D.D., Paul, S.M., Mennerick, S., and Holtzman, D.M. (2005). Synaptic activity regulates interstitial fluid amyloid-beta levels in vivo. Neuron 48, 913–922. 10.1016/j.neuron.2005.10.028.

84. Buxbaum, J.D., Thinakaran, G., Koliatsos, V., O’Callahan, J., Slunt, H.H., Price, D.L., and Sisodia, S.S. (1998). Alzheimer amyloid protein precursor in the rat hippocampus: transport and processing through the perforant path. J Neurosci 18, 9629–9637. 10.1523/JNEUROSCI.18-23-09629.1998.

85. Nitsch, R.M., Farber, S.A., Growdon, J.H., and Wurtman, R.J. (1993). Release of amyloid beta-protein precursor derivatives by electrical depolarization of rat hippocampal slices. Proc Natl Acad Sci U S A 90, 5191–5193. 10.1073/pnas.90.11.5191.

86. Groemer, T.W., Thiel, C.S., Holt, M., Riedel, D., Hua, Y., Huve, J., Wilhelm, B.G., and Klingauf, J. (2011). Amyloid precursor protein is trafficked and secreted via synaptic vesicles. PLoS One 6, e18754. 10.1371/journal.pone.0018754.

87. Carlson, G.A., and Prusiner, S.B. (2021). How an Infection of Sheep Revealed Prion Mechanisms in Alzheimer’s Disease and Other Neurodegenerative Disorders. Int J Mol Sci 22. 10.3390/ijms22094861.

88. Zarow, C., Lyness, S.A., Mortimer, J.A., and Chui, H.C. (2003). Neuronal loss is greater in the locus coeruleus than nucleus basalis and substantia nigra in Alzheimer and Parkinson diseases. Arch Neurol 60, 337–341. 10.1001/archneur.60.3.337.

89. Anuncibay Soto, B., Ma, Y., Nollet, M., Wong, S., Miracca, G., Rastinejad, D., Yustos, R., Vyssotski, A.L., Franks, N.P., and Wisden, W. (2025). The locus coeruleus maintains core body temperature and protects against hypothermia during dexmedetomidine-induced sedation. Proc Natl Acad Sci U S A 122, e2422878122. 10.1073/pnas.2422878122.

90. Arnsten, A.F., and Goldman-Rakic, P.S. (1985). Alpha 2-adrenergic mechanisms in prefrontal cortex associated with cognitive decline in aged nonhuman primates. Science 230, 1273–1276. 10.1126/science.2999977.

91. Yamada, M., Lamping, K.G., Duttaroy, A., Zhang, W., Cui, Y., Bymaster, F.P., McKinzie, D.L., Felder, C.C., Deng, C.X., Faraci, F.M., and Wess, J. (2001). Cholinergic dilation of cerebral blood vessels is abolished in M(5) muscarinic acetylcholine receptor knockout mice. Proc Natl Acad Sci U S A 98, 14096–14101. 10.1073/pnas.251542998.

92. Greenberg, S.M., Bacskai, B.J., Hernandez-Guillamon, M., Pruzin, J., Sperling, R., and van Veluw, S.J. (2020). Cerebral amyloid angiopathy and Alzheimer disease - one peptide, two pathways. Nat Rev Neurol 16, 30–42. 10.1038/s41582-019-0281-2.

93. Nizari, S., Wells, J.A., Carare, R.O., Romero, I.A., and Hawkes, C.A. (2021). Loss of cholinergic innervation differentially affects eNOS-mediated blood flow, drainage of Abeta and cerebral amyloid angiopathy in the cortex and hippocampus of adult mice. Acta Neuropathol Commun 9, 12. 10.1186/s40478-020-01108-z.

94. Nizari, S., Carare, R.O., Romero, I.A., and Hawkes, C.A. (2019). 3D Reconstruction of the Neurovascular Unit Reveals Differential Loss of Cholinergic Innervation in the Cortex and Hippocampus of the Adult Mouse Brain. Front Aging Neurosci 11, 172. 10.3389/fnagi.2019.00172.

95. Lee, M.G., Hassani, O.K., Alonso, A., and Jones, B.E. (2005). Cholinergic basal forebrain neurons burst with theta during waking and paradoxical sleep. J Neurosci 25, 4365–4369. 10.1523/JNEUROSCI.0178-05.2005.

96. Ma, X., Zhang, Y., Wang, L., Li, N., Barkai, E., Zhang, X., Lin, L., and Xu, J. (2020). The Firing of Theta State-Related Septal Cholinergic Neurons Disrupt Hippocampal Ripple Oscillations via Muscarinic Receptors. J Neurosci 40, 3591–3603. 10.1523/JNEUROSCI.1568-19.2020.

97. Peever, J., and Fuller, P.M. (2017). The Biology of REM Sleep. Curr Biol 27, R1237–R1248. 10.1016/j.cub.2017.10.026.

98. Kashiwagi, M., Beck, G., Kanuka, M., Arai, Y., Tanaka, K., Tatsuzawa, C., Koga, Y., Saito, Y.C., Takagi, M., Oishi, Y., et al. (2024). A pontine-medullary loop crucial for REM sleep and its deficit in Parkinson’s disease. Cell 187, 6272–6289 e6221. 10.1016/j.cell.2024.08.046.

99. Luppi, P.H., Malcey, J., Chancel, A., Duval, B., Cabrera, S., and Fort, P. (2025). Neuronal network controlling REM sleep. J Sleep Res 34, e14266. 10.1111/jsr.14266.

100. Miracca, G., Anuncibay-Soto, B., Tossell, K., Yustos, R., Vyssotski, A.L., Franks, N.P., and Wisden, W. (2022). NMDA Receptors in the Lateral Preoptic Hypothalamus Are Essential for Sustaining NREM and REM Sleep. J Neurosci 42, 5389–5409. 10.1523/JNEUROSCI.0350-21.2022.

101. Adamantidis, A.R., and de Lecea, L. (2023). Sleep and the hypothalamus. Science 382, 405–412. 10.1126/science.adh8285.

102. Wang, Z.Y., Fei, X., Liu, X.T., Wang, Y.J., Hu, Y., Peng, W.L., Wang, Y.W., Zhang, S.Y., and Xu, M. (2022). REM sleep is associated with distinct global cortical dynamics and controlled by occipital cortex. Nature Communications 13. ARTN 6896 10.1038/s41467-022-34720-9.

103. Dong, Y.F., Li, J.Q., Zhou, M., Du, Y.H., and Liu, D.Q. (2022). Cortical regulation of two-stage rapid eye movement sleep. Nature Neuroscience 25, 1675-+. 10.1038/s41593-022-01195-2.

104. Hong, J., Lozano, D.E., Beier, K.T., Chung, S., and Weber, F. (2023). Prefrontal cortical regulation of REM sleep. Nature Neuroscience 26, 1820-+. 10.1038/s41593-023-01398-1.

105. Hasan, S., Dauvilliers, Y., Mongrain, V., Franken, P., and Tafti, M. (2012). Age-related changes in sleep in inbred mice are genotype dependent. Neurobiol Aging 33, 195 e113–126. 10.1016/j.neurobiolaging.2010.05.010.

106. Rawlins, J.N., Feldon, J., and Gray, J.A. (1979). Septo-hippocampal connections and the hippocampal theta rhythm. Exp Brain Res 37, 49–63. 10.1007/BF01474253.

107. Calafate, S., Ozturan, G., Thrupp, N., Vanderlinden, J., Santa-Marinha, L., Morais-Ribeiro, R., Ruggiero, A., Bozic, I., Rusterholz, T., Lorente-Echeverria, B., et al. (2023). Early alterations in the MCH system link aberrant neuronal activity and sleep disturbances in a mouse model of Alzheimer’s disease. Nat Neurosci 26, 1021–1031. 10.1038/s41593-023-01325-4.

108. Drew, V.J., Wang, C., and Kim, T. (2023). Progressive sleep disturbance in various transgenic mouse models of Alzheimer’s disease. Front Aging Neurosci 15, 1119810. 10.3389/fnagi.2023.1119810.

109. Petit, D., Montplaisir, J., Lorrain, D., and Gauthier, S. (1992). Spectral analysis of the rapid eye movement sleep electroencephalogram in right and left temporal regions: a biological marker of Alzheimer’s disease. Ann Neurol 32, 172–176. 10.1002/ana.410320208.

110. Petit, D., Lorrain, D., Gauthier, S., and Montplaisir, J. (1993). Regional spectral analysis of the REM sleep EEG in mild to moderate Alzheimer’s disease. Neurobiol Aging 14, 141–145. 10.1016/0197-4580(93)90089-t.

111. Brayet, P., Petit, D., Frauscher, B., Gagnon, J.F., Gosselin, N., Gagnon, K., Rouleau, I., and Montplaisir, J. (2016). Quantitative EEG of Rapid-Eye-Movement Sleep: A Marker of Amnestic Mild Cognitive Impairment. Clin EEG Neurosci 47, 134–141. 10.1177/1550059415603050.

112. D’Atri, A., Scarpelli, S., Gorgoni, M., Truglia, I., Lauri, G., Cordone, S., Ferrara, M., Marra, C., Rossini, P.M., and De Gennaro, L. (2021). EEG alterations during wake and sleep in mild cognitive impairment and Alzheimer’s disease. iScience 24, 102386. 10.1016/j.isci.2021.102386.

113. D’Rozario, A.L., Chapman, J.L., Phillips, C.L., Palmer, J.R., Hoyos, C.M., Mowszowski, L., Duffy, S.L., Marshall, N.S., Benca, R., Mander, B., et al. (2020). Objective measurement of sleep in mild cognitive impairment: A systematic review and meta-analysis. Sleep Med Rev 52, 101308. 10.1016/j.smrv.2020.101308.

114. Martinez-Canada, P., Perez-Valero, E., Minguillon, J., Pelayo, F., Lopez-Gordo, M.A., and Morillas, C. (2023). Combining aperiodic 1/f slopes and brain simulation: An EEG/MEG proxy marker of excitation/inhibition imbalance in Alzheimer’s disease. Alzheimers Dement (Amst) 15, e12477. 10.1002/dad2.12477.

115. Eyamu, J., Ku, B., Kim, K., Lee, K.H., and Kim, J.U. (2026). Aperiodic EEG signatures: unveiling the interplay between APOE ε4 and mild cognitive impairment subtypes. Frontiers in Aging Neuroscience 17. ARTN 1675330 10.3389/fnagi.2025.1675330.

116. Nanclares, C., Colmena, I., Munoz-Montero, A., Baraibar, A.M., de Pascual, R., Wojnicz, A., Ruiz-Nuno, A., Garcia, A.G., Gironda-Martinez, A., and Gandia, L. (2025). Beyond the brain: early autonomic dysfunction in Alzheimer’s disease. Acta Neuropathol Commun 13, 128. 10.1186/s40478-025-02042-8.

117. Okada, K., Hashimoto, K., and Kobayashi, K. (2022). Cholinergic regulation of object recognition memory. Front Behav Neurosci 16, 996089. 10.3389/fnbeh.2022.996089.

118. Solari, N., and Hangya, B. (2018). Cholinergic modulation of spatial learning, memory and navigation. Eur J Neurosci 48, 2199–2230. 10.1111/ejn.14089.

119. Wu, J.L., Li, Z.M., Chen, H., Chen, W.J., Hu, N.Y., Jin, S.Y., Li, X.W., Chen, Y.H., Yang, J.M., and Gao, T.M. (2024). Distinct septo-hippocampal cholinergic projections separately mediate stress-induced emotional and cognitive deficits. Sci Adv 10, eado1508. 10.1126/sciadv.ado1508.

120. Shivakumar, A.B., Mehak, S.F., Gupta, A., and Gangadharan, G. (2025). Medial septal cholinergic neurotransmission is essential for social memory in mice. Prog Neuropsychopharmacol Biol Psychiatry 136, 111207. 10.1016/j.pnpbp.2024.111207.

121. Pimpinella, D., Mastrorilli, V., Giorgi, C., Coemans, S., Lecca, S., Lalive, A.L., Ostermann, H., Fuchs, E.C., Monyer, H., Mele, A., et al. (2021). Septal cholinergic input to CA2 hippocampal region controls social novelty discrimination via nicotinic receptor-mediated disinhibition. Elife 10. 10.7554/eLife.65580.

122. Chung, J.A., and Cummings, J.L. (2000). Neurobehavioral and neuropsychiatric symptoms in Alzheimer’s disease: characteristics and treatment. Neurol Clin 18, 829–846. 10.1016/s0733-8619(05)70228-0.

123. Bellio, T.A., Laguna-Torres, J.Y., Campion, M.S., Chou, J., Yee, S., Blusztajn, J.K., and Mellott, T.J. (2024). Perinatal choline supplementation prevents learning and memory deficits and reduces brain amyloid Abeta42 deposition in AppNL-G-F Alzheimer’s disease model mice. PLoS One 19, e0297289. 10.1371/journal.pone.0297289.

124. Cavedo, E., Grothe, M.J., Colliot, O., Lista, S., Chupin, M., Dormont, D., Houot, M., Lehericy, S., Teipel, S., Dubois, B., et al. (2017). Reduced basal forebrain atrophy progression in a randomized Donepezil trial in prodromal Alzheimer’s disease. Sci Rep 7, 11706. 10.1038/s41598-017-09780-3.

125. Rossi, J., Balthasar, N., Olson, D., Scott, M., Berglund, E., Lee, C.E., Choi, M.J., Lauzon, D., Lowell, B.B., and Elmquist, J.K. (2011). Melanocortin-4 receptors expressed by cholinergic neurons regulate energy balance and glucose homeostasis. Cell Metab 13, 195–204. 10.1016/j.cmet.2011.01.010.

126. Yang, C.F., Chiang, M.C., Gray, D.C., Prabhakaran, M., Alvarado, M., Juntti, S.A., Unger, E.K., Wells, J.A., and Shah, N.M. (2013). Sexually dimorphic neurons in the ventromedial hypothalamus govern mating in both sexes and aggression in males. Cell 153, 896–909. 10.1016/j.cell.2013.04.017.

127. Yu, X., Li, W., Ma, Y., Tossell, K., Harris, J.J., Harding, E.C., Ba, W., Miracca, G., Wang, D., Li, L., et al. (2019). GABA and glutamate neurons in the VTA regulate sleep and wakefulness. Nat Neurosci 22, 106–119. 10.1038/s41593-018-0288-9.

128. Klugmann, M., Symes, C.W., Leichtlein, C.B., Klaussner, B.K., Dunning, J., Fong, D., Young, D., and During, M.J. (2005). AAV-mediated hippocampal expression of short and long Homer 1 proteins differentially affect cognition and seizure activity in adult rats. Mol Cell Neurosci 28, 347–360. 10.1016/j.mcn.2004.10.002.

129. Anisimov, V.N., Herbst, J.A., Abramchuk, A.N., Latanov, A.V., Hahnloser, R.H., and Vyssotski, A.L. (2014). Reconstruction of vocal interactions in a group of small songbirds. Nat Methods 11, 1135–1137. 10.1038/nmeth.3114.

130. Gunaydin, L.A., Grosenick, L., Finkelstein, J.C., Kauvar, I.V., Fenno, L.E., Adhikari, A., Lammel, S., Mirzabekov, J.J., Airan, R.D., Zalocusky, K.A., et al. (2014). Natural neural projection dynamics underlying social behavior. Cell 157, 1535–1551. 10.1016/j.cell.2014.05.017.

131. Kraeuter, A.K., Guest, P.C., and Sarnyai, Z. (2019). The Y-Maze for Assessment of Spatial Working and Reference Memory in Mice. Methods Mol Biol 1916, 105–111. 10.1007/978-1-4939-8994-2_10.

132. Leger, M., Quiedeville, A., Bouet, V., Haelewyn, B., Boulouard, M., Schumann-Bard, P., and Freret, T. (2013). Object recognition test in mice. Nat Protoc 8, 2531–2537. 10.1038/nprot.2013.155.

133. Matikainen-Ankney, B.A., Garmendia-Cedillos, M., Ali, M., Krynitsky, J., Salem, G., Miyazaki, N.L., Pohida, T., and Kravitz, A.V. (2019). Rodent Activity Detector (RAD), an Open Source Device for Measuring Activity in Rodent Home Cages. eNeuro 6. 10.1523/ENEURO.0160-19.2019.

134. Abelson, K.S., Kalliokoski, O., Teilmann, A.C., and Hau, J. (2016). Applicability of Commercially Available ELISA Kits for the Quantification of Faecal Immunoreactive Corticosterone Metabolites in Mice. In Vivo 30, 739–744. 10.21873/invivo.10989.

